# Learning Strand-Equivariant Tokenizations of DNA with MirrorDNA

**DOI:** 10.64898/2026.11.04.764207

**Authors:** Nirjhor Datta, Swakkhar Shatabda, Md. Shamsuzzoha Bayzid, M. Sohel Rahman

## Abstract

DNA has two strands, and for most regulatory label prediction tasks a sequence and its reverse complement should get same answer. Yet DNA language models are tokenized, trained and tested on one strand at a time. We show that this makes them inconsistent. BPE and *k*-mer tokenizers split 92–100% of transcription-factor motifs differently on the two strands, and NT-v2, DNABERT-2 and HyenaDNA change 25-31% of their predictions when the test sequences are reverse-complemented. We introduce MirrorDNA, a compact DNA language model with a learned tokenizer that guarantees mirror-symmetric token boundaries across reverse-complement sequences, regardless of its learned parameters. We prove that this symmetry holds through the whole model. As a result, the same model can give identical answers on both strands when the label does not depend on the strand, and can still tell the strands apart when it does, which no strand-blind model can learn. Counting only predictions that are correct on both strands, MirrorDNA more than doubles the MCC of NT-v2, DNABERT-2 and HyenaDNA. With 2.3M parameters, it is more accurate than the Caduceus models trained on the same data, and not significantly different from NT-v2 (98M parameters) on Genomic Benchmarks or from DNABERT-2 (117M) on the Nucleotide Transformer tasks. It also stays accurate on inputs sixteen times longer than those it was trained on. The same model trained with reverse-complement augmentation instead still changes about one prediction in ten with the strand, unless its outputs are averaged over both strands. Without any annotation, the tokenizer learns to keep the splice-donor signal `AGGT` in a single token. Code: https://github.com/NIRJHOR-DATTA/MirrorDNA.

## 1 Introduction

Language models pretrained on genomic sequence are now a common starting point in regulatory genomics. Examples include DNABERT [18], the Nucleotide Transformer [9], DNABERT-2 [36], GROVER [28], HyenaDNA [25],

Caduceus [29], GPN [4, 3] and Evo [24, 6], which follow supervised networks such as DeepSEA and Enformer [1, 35, 20, 19, 17, 2]. Large models are expensive to adapt. Frozen embeddings can match fine-tuning at 10–20× lower cost [10], but position-precise signal, for instance at splice sites, is often lost in pooled frozen representations [11]. How much large-scale pretraining adds is still debated [33], and compact convolutional models remain competitive [5]. We therefore focus on small models fine-tuned end to end.

Every one of these models has to decide how DNA is cut into tokens. Single-nucleotide tokens keep full resolution but make sequences long [25, 29]. *k*-mers [18, 9] and byte-pair encoding (BPE) [30, 36] shorten the input with a vocabulary fixed before training, and the choice of vocabulary changes downstream accuracy [13]. BiRNA-BERT [32] combines nucleotide and BPE tokens for RNA. More recent work learns the segmentation together with the model. MxDNA [27] and VQDNA [22] learn a vocabulary, while dynamic chunking [16], patch-based byte models [26] and DNAChunker [21] place boundaries according to content. This suits genomes well, because motifs, splice signals and repeats have no fixed length.

DNA has a symmetry that text lacks. Each locus can be read on either of two complementary strands, and for many labels (histone marks, chromatin accessibility, most enhancer annotations) the orientation of the reference sequence means nothing. A model that predicts differently for a sequence and its reverse complement is then simply inconsistent with the biology. Other labels do depend on the strand. A promoter points towards its gene, splice sites are defined by the direction of transcription, and a gene lies on one strand or the other. A useful model should agree with itself across strands when the biology is symmetric, and tell the strands apart when it is not.

Current methods handle only one side of this. Reverse-complement augmentation [19, 29] encourages consistency without guaranteeing it, and averaging both strands [35] doubles the inference cost. Parameter sharing [31] and equivariant layers [23, 34, 8, 7] build the symmetry into convolutional models. Caduceus [29] brought it to pretrained language models with an equivariant Mamba [15] backbone, but its standard classification head averages the two strands and so cannot predict a label that changes with the strand. All of these models work on single nucleotides. Learned tokenizers, for their part, ignore the symmetry. A boundary predictor that does not respect it cuts the two strands at unrelated positions, so the two strands get different tokens and later layers are no longer guaranteed to agree. Neither MxDNA nor DNAChunker addresses the reverse complement [27, 21].

The problem is not a corner case. Take a transcription-factor binding site and read it on each strand. BPE (as in DNABERT-2) splits it into different tokens on the two strands in 92% of cases, and non-overlapping 6-mers (as in the Nucleotide Transformer) in 100%. A learned chunker trained with reverse-complement augmentation still does so in 88%. Standard benchmarks [9, 14] test each window in one orientation only. If we reverse-complement the test windows of strand-symmetric tasks, NT-v2, DNABERT-2 and HyenaDNA change their answer for 25–31% of sequences. Counting only predictions that are right on both strands, their MCC drops to 0.08-0.19, from 0.51-0.54 on the forward strand.

We present MirrorDNA, a small DNA language model with a learned tokenizer that treats both strands the same way by design. Whatever its parameters, it cuts the two strands of a sequence into mirror-image chunks. Our contributions are threefold.

1. **A strand-equivariant learned tokenizer, with guarantees**. A symmetric boundary router cuts the two strands at mirrored positions, and a shared-weight encoder reads both strands of every base pair. We prove that the whole model, from segmentation to output, is equivariant to the reverse complement. A model that always gives the same answer on both strands cannot learn labels that depend on the strand. The backbone therefore carries two heads, one that gives the same answer on both strands and one that can tell them apart (Section 2). To our knowledge, MirrorDNA is the first learned DNA tokenizer with this property.
2. **Strand-aware evaluation**. We use all 18 tasks of the revised Nucleotide Transformer benchmark [9], all 9 Genomic Benchmarks [14] and two gene-strand tasks built from RefSeq. We report accuracy on each strand, a *both-strands* MCC that counts a prediction as correct only if it is correct on both strands, and the share of predictions that change with the strand. All results are over three seeds, with paired tests.
3. **Consistent, accurate and label-efficient at small scale**. With 2.3M parameters, and the same pretraining data and token budget as the 1-kb Caduceus models [29], MirrorDNA gives the same answer on both strands and reaches a both-strands MCC of 0.491 ± 0.004. That is more than twice the value of NT-v2, DNABERT-2 and HyenaDNA, and higher on all 12 strand-symmetric tasks (Wilcoxon *p <* 0.001).It is more accurate than the 1-kb Caduceus models over all 29 tasks (*p <* 0.001), and not significantly different from the 98M-parameter NT-v2 on Genomic Benchmarks (0.751 vs. 0.741, *p* = 0.36) or from the 117M-parameter DNABERT-2 on the Nucleotide Transformer tasks (0.618 vs. 0.631, *p* = 0.42). We also compare built-in symmetry with reverse-complement augmentation in the same architecture. With all labels the two are not significantly different. With 2,000 labels per task, the equivariant model is more accurate than one pass of the augmented model ( +0.018 MCC, *p <* 0.001), which still changes 10–12% of its predictions with the strand. Averaging the augmented model over both strands closes this gap. It keeps its accuracy on inputs sixteen times longer than in pretraining. Without any annotation, its tokenizer turns the splice-donor signal `AGGT` into a single token nine times more often than expected by chance.

## 2 Methods

MirrorDNA follows the layout of dynamic-chunking language models [16, 21] (Fig. 1). An encoder produces one vector per base, a router decides where to cut, the chunks are pooled and passed through a transformer, and a light decoder maps the outputs back to bases for masked-language-model pretraining. In MirrorDNA, every stage, including the choice of chunk boundaries, commutes with the reverse complement for any parameter values, so the property needs no augmentation and cannot be lost in training.

**Fig. 1.**
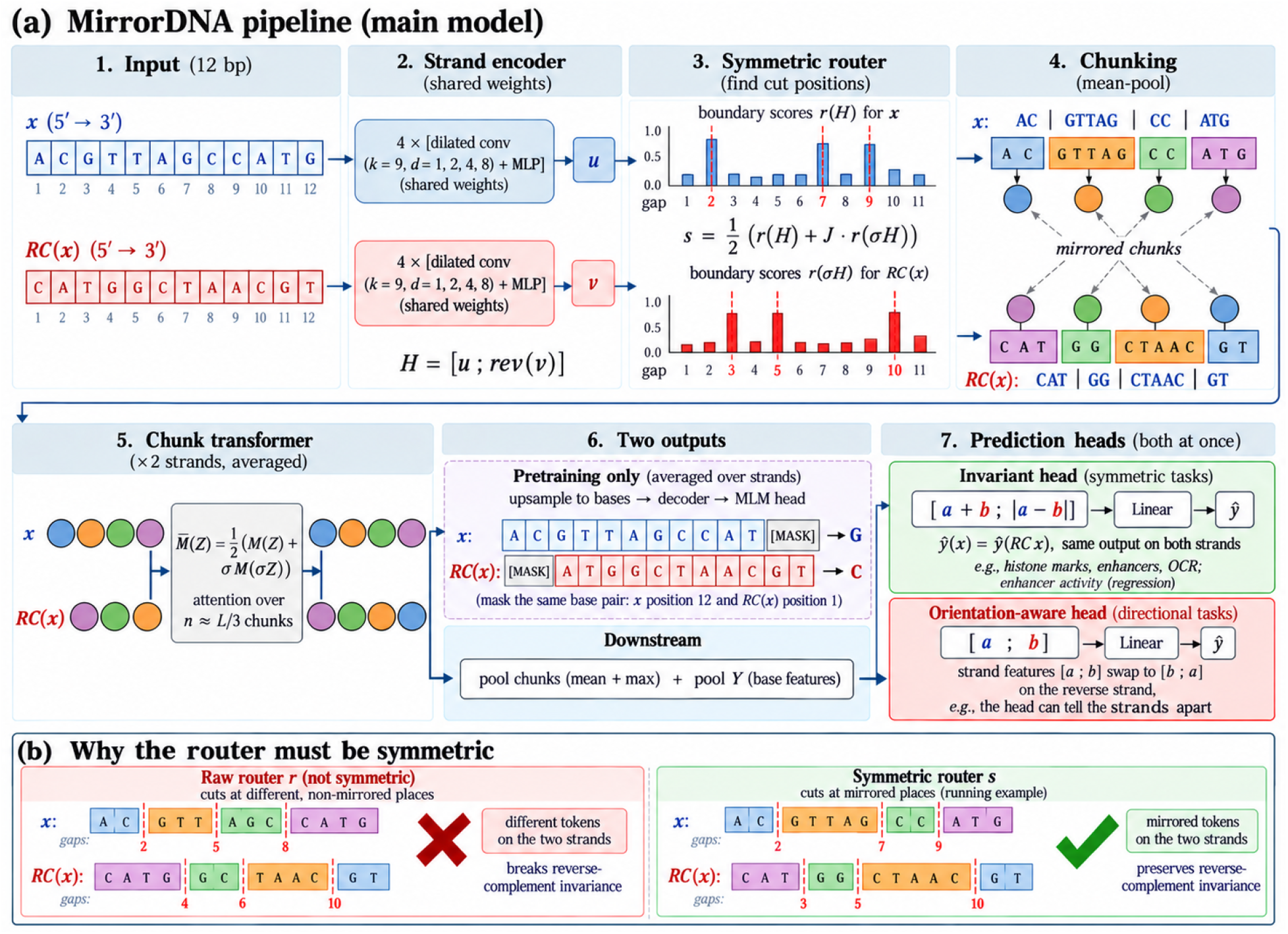
Overview of MirrorDNA. (a) Both strands are encoded with shared weights and cut at mirrored positions. Chunks are pooled and processed by a group-averaged transformer, and two heads serve strand-symmetric and direction-dependent tasks. (b) A raw router cuts the two strands differently. The symmetric router cuts them at mirrored positions, so each chunk on one strand is the reverse complement of a chunk on the other.

### Notation

For *x* ∈ {A, C, G, T}^*L*^ with complement *c*, the reverse complement is ℛ (*x*)_*i*_ = *c*(*x*_*L*+1−*i*_). Hidden states *H* ∈ ℝ^*L*×2*d*^ have two halves of width *d*, one per strand. The strand action *σ* (*H*)_*i*_ = *P H*_*L*+1−*i*_ reverses the positions and swaps the two halves (*P* [*u*; *v*] = [*v*; *u*]). On *n* chunk vectors it is *σ*_*n*_ (*Z*)_*k*_ = *P Z*_*n*+1−*k*_, and on the *L* − 1 gaps between adjacent bases it is the reversal (*J p*) _*j*_ = *p*_*L*− *j*_ . We call a map *F strand-equivariant* if *F* (ℛ*x*) = *σF* (*x*), and *strand-invariant* if *F* (R*x*) = *F* (*x*).

### 2.1 Strand encoder and symmetric router

The encoder *g* has four residual blocks, each with layer normalization, a depthwise convolution (kernel 9, dilations 1, 2, 4 and 8) and an MLP. It reads *x* and ℛ (*x*) with the same weights, each in its own 5^′^ → 3^′^ direction, and we set *H (x )*= [*g x* ; rev *g* (*ℛ x*)] . Every base therefore carries the context of both strands. Swapping *x* and ℛ (*x)* swaps the two halves, so *H(*ℛ *x* = *σH (x )*(Supplementary Lemma S1).

As in H-Net [16], a boundary is likely where adjacent bases look different. The raw score of the gap between bases *j* and *j*+ 1 is 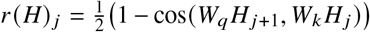 . This score is not mirror-symmetric, because it compares each base with its left neighbour and *W*_*q*_ ≠ *W*_*k*_ . Cutting the two strands with *r* therefore gives different chunks, and later layers are then not guaranteed to agree across strands. Instead, we cut gap *j* whenever the symmetrized score

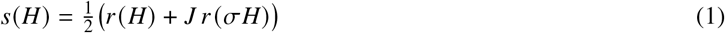

is at least 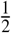 . Each chunk is represented by the mean of *H* over its bases, giving *Z* ∈ ℝ^*n*×2*d*^.

#### Lemma 1

**(Mirrored segmentation)**. *For any router r, s( σH)* = *J s (H)* . *Hence, since H(ℛ x)* = *σH (x), the cut sets of the two strands are mirror images,C (ℛx)* = {*L* − *j* : *j ∈ Cx)}, chunk k of ℛx is chunk n+* 1 −*k of x reverse-complemented, and Z (ℛx)* = *σ*_*n*_ *Z x* .

*Proof*. Since *σ*^2^ = *J*^2^ = id, we have 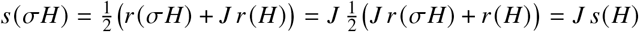. So gap *L* − *j* of ℛ*x* gets the score of gap *j* of *x*, and the same threshold rule gives the same decision. A mean does not depend on the order of its terms. □

The router is trained as in H-Net, through a straight-through chunk confidence and a ratio loss that keeps the fraction of cut gaps near 1/4. Both terms are mirror-symmetric (Supplementary Section S2).

### 2.2 Chunk transformer, decoder and pretraining

A pre-norm transformer *M* with sinusoidal positions processes the *n* chunks, of which there are about one per three bases after training. A transformer is not equivariant to *σ*_*n*_, so we use its group average 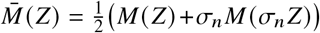 which satisfies 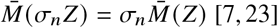. Like Caduceus-PS, the model thus sees both strands of every input. Each base receives the output of its chunk, scaled by the chunk confidence (*U*). A decoder *D*, made of a residual projection of *H* followed by two convolutional blocks, is group-averaged as a whole, 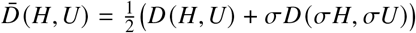, so the per-base states 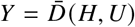 satisfy *Y* (ℛ*x*) = *σY* (*x*). Finally, the masked-LM head *h* is averaged with its own prediction on the other strand, 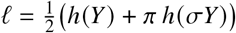, where *π* reverses positions and complements bases.

We pretrain with the 1-kb recipe of Caduceus and HyenaDNA [29, 25], on 1,024-bp windows from the Basenji/Enformer intervals of hg38 [19, 2], with 2^20^ bases per step for 10,000 steps (10.5B bases) and 15% of bases masked. The main model has *d* = 96, four transformer layers and 2.3M parameters. We also release MirrorDNA-duplex (2.5M), whose encoder gates each strand with the other at the same base pair. It shares its architecture with the non-equivariant control below, which isolates the effect of the symmetry.

### 2.3 Downstream heads and guarantees

Let [*a*; *b* ]denote the two strand halves of the mean (or max) of 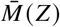 over chunks. For strand-symmetric tasks, the *invariant head* uses [*a+ b*,| *a* −*b*|] . For labels defined relative to the strand being read (promoters, splice sites, gene orientation), the *orientation-aware head* keeps *a*; *b* as it is. Both heads are linear layers that output class logits, or one value per target for regression, and both also receive the same pooled combinations of *Y* . A *centre readout* can add *Y* pooled in the same way over the central 8, 32 and 128 bases of the window, where the signal of site-centred tasks such as splice sites lies. The central window of ℛ*x* is the central window of *x* read backwards, so these features obey the same identities and Theorem 1 holds unchanged.

#### Theorem 1

**(End-to-end equivariance; proof in Supplementary Section S1)**. *For every parameter value and input x (the model has no dropout), l (ℛx)* = *π l (x) and [a*; *b*]*( ℛ x)* = [*b*; *a*] *(x)* . *Hence the invariant features, and any classifier or regressor built on them, give identical outputs on both strands, while the orientation-aware features are swapped*.

For strand-symmetric labels and convex losses, an invariant Bayes-optimal predictor exists (Supplementary Proposition S1), so requiring invariance does not exclude the best predictor, although our features need not keep every invariant distinction. When the reverse complement of one class is the other class, however, invariance rules out learning altogether.

#### Proposition 1

**(Invariant heads cannot read orientation)**. *If X, Y and (ℛX*, 1 *Y) have the same distribution, with Y ∈ {*0, 1}, *then every strand-invariant classifier f has accuracy exactly*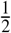, *and f (X) is independent of Y* . *Its population MCC is therefore zero whenever it is defined (that is, unless f is constant)*.

*Proof*. By the symmetry and the invariance of *f*, Pr( *f* ( *X*) = *Y* ) = Pr( *f* (ℛ *X*) = 1 − *Y* ) = Pr( *f* ( *X*) = 1 − *Y* ) = 1 − Pr( *f* ( *X*) = *Y* ). In the same way, Pr( *f* ( *X*) = 1, *Y* = 1) = Pr( *f* ( *X*) = 1, *Y* = 0), which gives 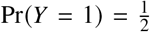 and independence. □

### 2.4 Tasks, baselines and evaluation

#### Tasks

We use all 18 tasks of the revised Nucleotide Transformer (NT) benchmark [9], of which 12 are strand-symmetric (histone marks, enhancers) and 6 direction-dependent (promoters, splice sites), and all 9 Genomic Benchmarks [14], each with its official split. We add two *gene-strand* tasks built from RefSeq (hg38). Each input is a 1-kb window centred on a transcription start site or a splice donor, and the model must tell whether the gene lies on the + or the− strand. Chromosomes 8 and 9 are held out for testing. Because genes on both strands are sampled from the same genome, these tasks satisfy the condition of Proposition 1 approximately, though not by construction. We repeat them with 2-to 16-kb windows to test longer inputs. For regression we use DeepSTARR [12], which measures developmental and housekeeping enhancer activity for 249-bp *Drosophila* sequences (official split, scored by Pearson’s *r*).

#### Baselines

We compare with NT-v2-100M [9], DNABERT-2 [36], HyenaDNA-small [25] and the 1-kb Caduceus checkpoints [29], which were pretrained on the same data with the same token budget as MirrorDNA. Caduceus-PS is used with its published strand-invariant head, and Caduceus-PH with its published protocol, that is, with reverse-complement augmentation during fine-tuning (except on gene strand). To isolate the effect of the symmetry, we also train a learned chunker with exactly the architecture and pretraining of MirrorDNA-duplex that is not symmetrized but trained with reverse-complement augmentation instead (*Chunker + RC aug*.). For every model we also report *test-time averaging*, the prediction averaged over a sequence and its reverse complement. Since averaging acts only on the output, we also test the pretrained representations themselves. Here every model is used frozen. On the 12 strand-symmetric NT tasks, we compare the pooled embedding of each test sequence with that of its reverse complement. We also train a logistic-regression probe on the forward embeddings (up to 20,000 sequences per task) and test it on both strands.

#### Fine-tuning and evaluation

All models are fine-tuned end to end in the same way. We use AdamW (learning rate 3×10^−4^, or 3× 10^−5^ for NT-v2 and DNABERT-2), at most 10 epochs with early stopping (patience 3) on a 10% validation split, and the same mean-and-max pooled linear head. In splice tasks such as splice sites the signal sits at the center of the window, and pooling over the whole window loses its position. MirrorDNA is therefore fine-tuned with both heads, and for each task we keep the one with the higher validation MCC (mean over seeds), so the test sets play no part in the choice. The centre readout is kept for 7 of the 29 tasks: the three splice-site tasks, both gene-strand tasks, mouse enhancers and Ensembl regulatory elements. On DeepSTARR, MirrorDNA uses the pooled head. All other models, including MirrorDNA-duplex and the augmented chunker, use the pooled head. For label efficiency, training sets are subsampled to 500 or 2,000 sequences per task (Supplementary Section S2). Each test set is evaluated as given and reverse-complemented. For strand-symmetric tasks we report the *flip rate*, the share of test sequences whose prediction changes with the strand, and the *both-strands* MCC. The both-strands MCC counts a test sequence as correct only if the predictions for it and for its reverse complement are both correct. Formally, the both-strands prediction is *ŷ* (*x)* if *ŷ* (*x)* = *ŷ* (ℛ*x)* ; otherwise it is whichever of the two predictions is wrong (the forward one if both are), and we compute its MCC against the label *y*. For DeepSTARR we report how well the predictions for the two strands correlate and how often they differ by more than half a standard deviation. All numbers are means and standard deviations over three seeds, and models are compared with a two-sided Wilcoxon signed-rank test over tasks.

#### Tokenization analyses

On held-out chromosomes 8 and 9 we compare MirrorDNA, the augmented chunker, BPE and 6-mers in two ways. First, we compute the Jaccard index of the two strands’ (mirrored) cut sets in 2,000 random 1-kb windows. Second, we count the JASPAR 2024 vertebrate motif occurrences (relative score ≥0.85) that are tokenized differently on the two strands. At RefSeq splice sites, we compare the chunk that starts the intron or exon with the chunk found after shifting the same cuts by 100–900 bp, which keeps chunk lengths but breaks any link to the site. Finally, we shift 1-kb windows around 2,000 RefSeq splice donors by 1–32 bp and compare their boundaries with those of the centred window in genome coordinates, excluding 128 bp at each window edge.

## 3 Results

### 3.1 Tokenizers and models break strand symmetry, MirrorDNA does not

We first looked at how tokenizers treat the two strands of the same DNA (Fig. 2). The chunker trained with reverse-complement augmentation cuts them at largely different places (Jaccard index 0.39, no window cut identically). It also tokenizes 88% of JASPAR motif occurrences differently on the two strands. For BPE the figure is 92%, and for 6-mers it is 100%. MirrorDNA cuts every window and every motif occurrence identically on both strands (Jaccard index 1.00, 0% of occurrences), as Lemma 1 requires. Consistency across strands is distinct from consistency across window positions. Shifting the window by 1–32 bp leaves the boundaries of BPE, of the augmented chunker and of MirrorDNA unchanged away from the window edges (Jaccard index 1.000), whereas non-overlapping 6-mers and a fixed grid of mirrored 3-bp cuts move with the window (0.000 and 0.078). MirrorDNA is the only tokenizer that is consistent under both.

**Fig. 2.**
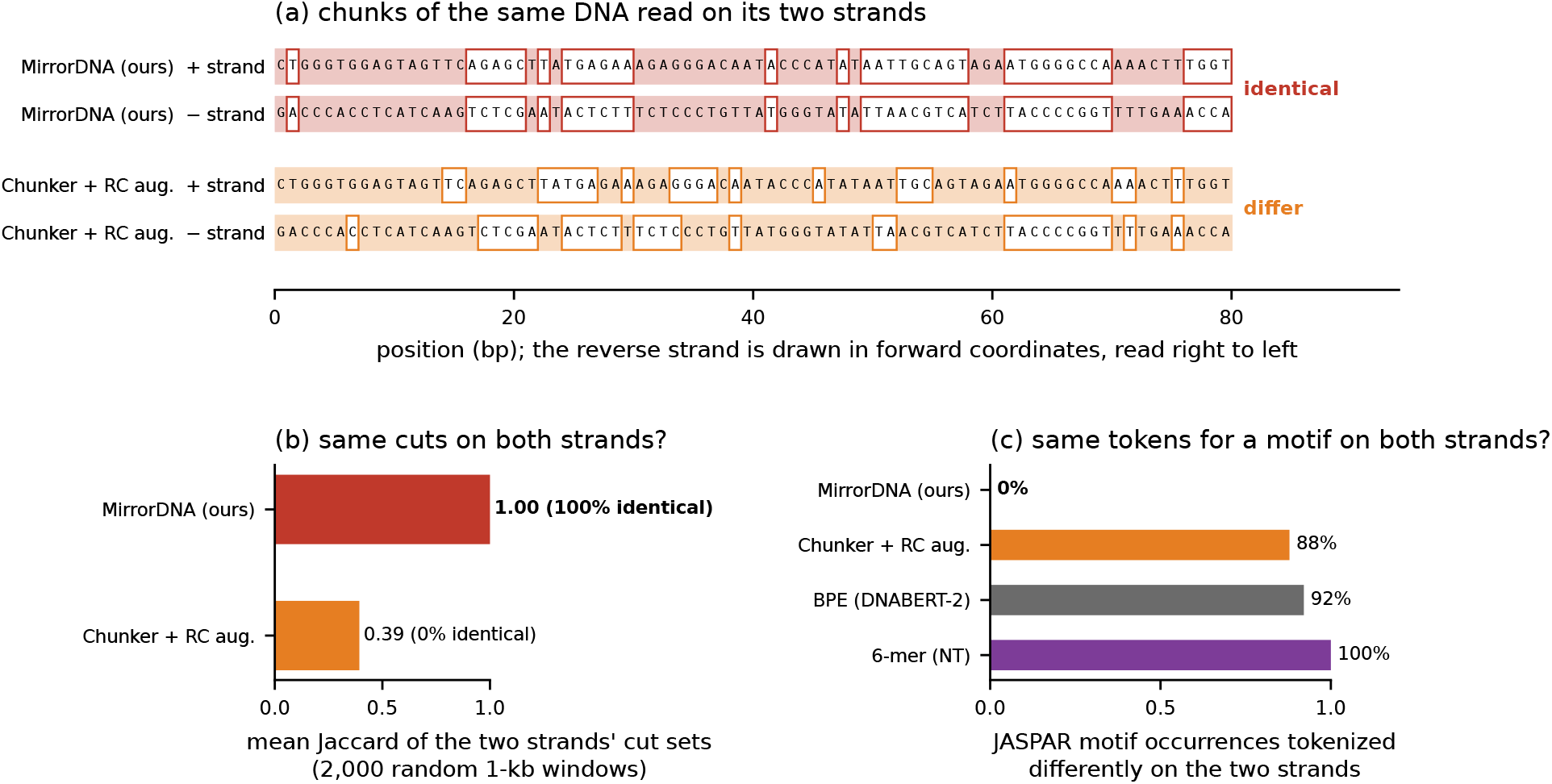
Tokenization of the two strands of the same DNA. (a) Chunks of 80 bp of a held-out window read on each strand. (b) Jaccard index of the two strands’ cut sets (2,000 random 1-kb windows). (c) JASPAR motif occurrences tokenized differently on the two strands.

These differences carry over to the predictions (Fig. 3, Table 1). On the 12 strand-symmetric NT tasks, NT-v2, DNABERT-2 and HyenaDNA change their prediction for 28.2%, 25.4% and 31.1% of the reverse-complemented test windows. If only predictions that are correct on both strands are counted, their MCC is 0.152, 0.187 and 0.080, compared with 0.51–0.54 on the forward strand. On the reverse strand alone, their MCC falls to 0.334, 0.350 and 0.272, whereas MirrorDNA gives 0.491 on both. On the 7 strand-symmetric Genomic Benchmarks tasks the effect is smaller (NT-v2: 0.710 forward, 0.685 RC, 11.1% flips). There MirrorDNA (0.711 on both strands) matches the forward MCC of NT-v2 and has the highest RC and both-strands MCC, ahead of Caduceus-PS-1k (0.686). With augmentation, Caduceus-PH-1k still flips 9.8% of its predictions and the augmented chunker 11.0% (both-strands MCC 0.389 and 0.381). MirrorDNA reaches a both-strands MCC of 0.491± 0.004. This is higher than every non-equivariant model on all 12 tasks (Wilcoxon *p <* 0.001 against each) and not significantly different from Caduceus-PS-1k, the other equivariant model (0.487, *p* = 0.15). The few flips we observe for MirrorDNA (0.003%) are floating-point ties at the decision threshold (Supplementary Section S2).

**Table 1.**
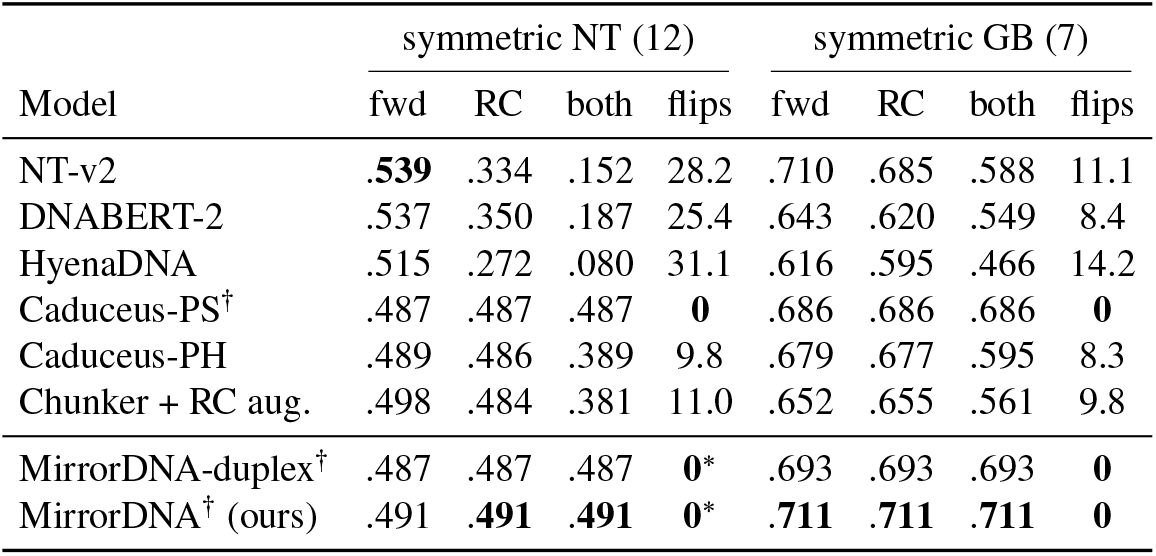
Strand consistency on the strand-symmetric tasks (12 NT, 7 Genomic Benchmarks; mean over 3 seeds). Each test set is evaluated as given (fwd) and reverse-complemented (RC); both: MCC of predictions correct on both strands; flips: % of predictions that change with the strand. Bold: best per column. MirrorDNA: head chosen per task on validation (Section 2.4). SDs: Supplementary Table S16. †Exactly equivariant. *Floating-point ties (0.003%).

| Model | symmetric NT (12) |  |  |  | symmetric GB (7) |  |  |  |
|---|---|---|---|---|---|---|---|---|
|  | fwd | RC | both | flips | fwd | RC | both | flips |
| NT-v2 | **.539** | .334 | .152 | 28.2 | .710 | .685 | .588 | 11.1 |
| DNABERT-2 | .537 | .350 | .187 | 25.4 | .643 | .620 | .549 | 8.4 |
| HyenaDNA | .515 | .272 | .080 | 31.1 | .616 | .595 | .466 | 14.2 |
| Caduceus-PS<sup>†</sup> | .487 | .487 | .487 | **0** | .686 | .686 | .686 | **0** |
| Caduceus-PH | .489 | .486 | .389 | 9.8 | .679 | .677 | .595 | 8.3 |
| Chunker + RC aug. | .498 | .484 | .381 | 11.0 | .652 | .655 | .561 | 9.8 |
| MirrorDNA-duplex<sup>†</sup> | .487 | .487 | .487 | **0**\* | .693 | .693 | .693 | **0** |
| MirrorDNA<sup>†</sup> (ours) | .491 | **.491** | **.491** | **0**\* | **.711** | **.711** | **.711** | **0** |

**Table 2.**
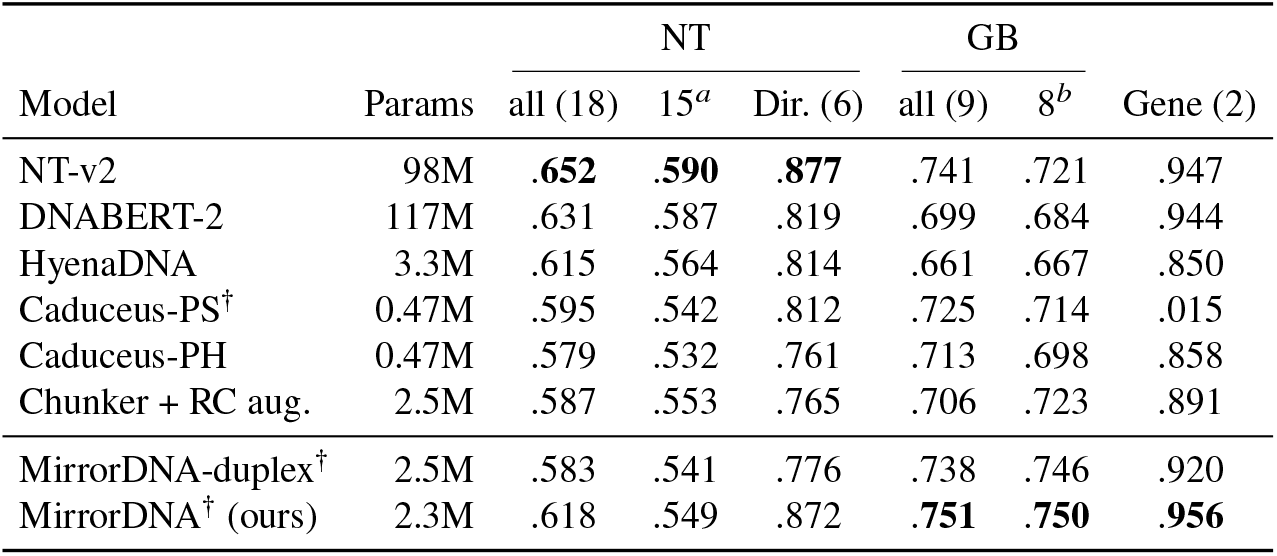
**Accuracy** (forward MCC, mean over 3 seeds) on the Nucleotide Transformer (NT) tasks, the Genomic Benchmarks (GB) and the 2 gene-strand tasks (Gene). Dir. = the 6 direction-dependent NT tasks. *a*NT without the 3 splice-site tasks, which depend on single-base precision at the junction. *b*GB without Ensembl regulatory elements, on which all three chunk-based models trail the single-nucleotide models. MirrorDNA: head chosen per task on validation (Section 2.4). Bold: best per column. Caduceus: the 1-kb checkpoints. SDs: Supplementary Table S13; per-task results: Supplementary Section S3. †Exactly equivariant.

**Fig. 3.**
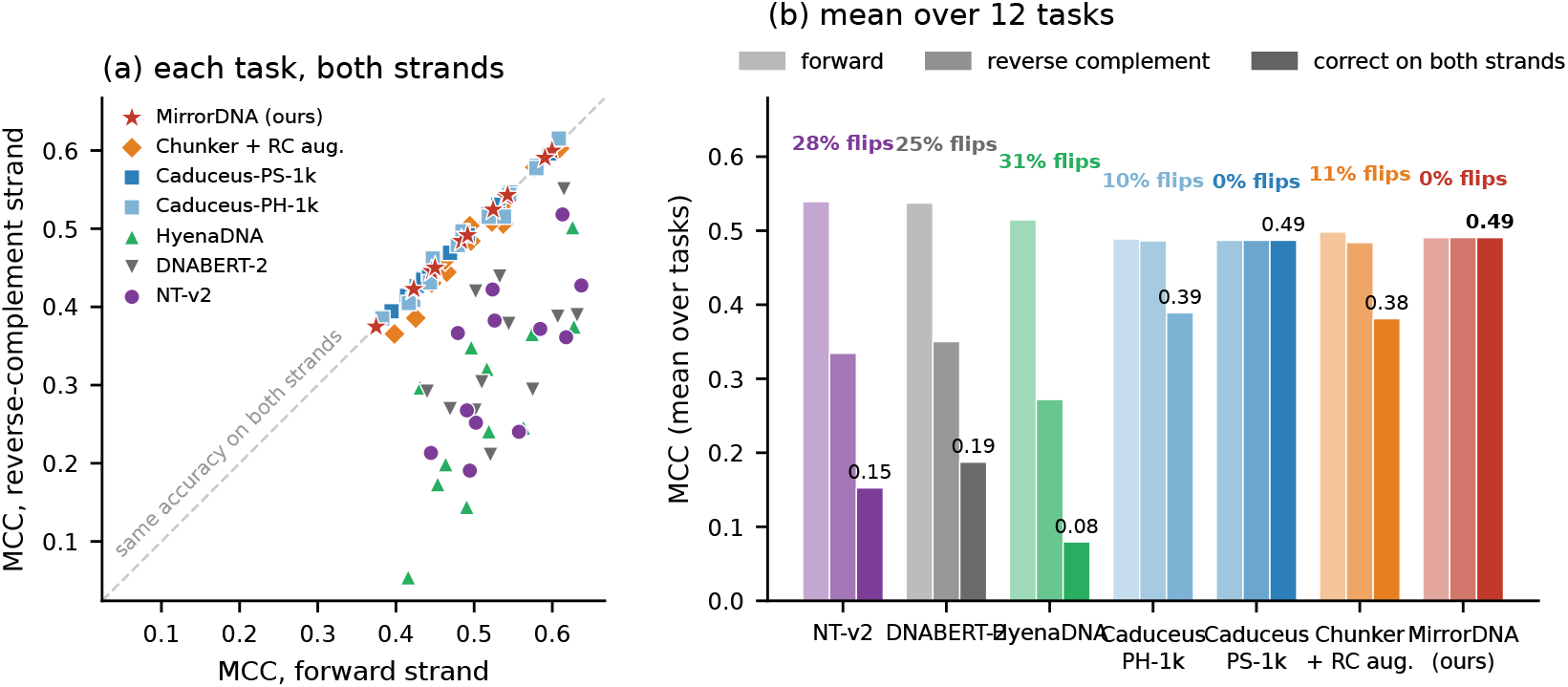
Accuracy on both strands. (12 strand-symmetric NT tasks, mean of 3 seeds). (a) Forward against reverse-complement MCC per task. (b) Forward, reverse-complement and both-strands MCC, and flip rates.

#### Regression

We see the same pattern for enhancer activity on DeepSTARR (Supplementary Section S3.4). For the two augmented models, the predictions for a sequence and for its reverse complement correlate at only 0.908 (Caduceus-PH-1k) and 0.901 (augmented chunker). They differ by more than half a standard deviation for 10.8% and 12.1% of sequences. The equivariant models predict the same activity on both strands. Mean Pearson’s *r* is similar across models (0.705 for MirrorDNA, 0.715 and 0.721 for Caduceus-PS/PH-1k) and close to the published DeepSTARR network (0.71) [12]. Averaging the predictions of the augmented models over both strands makes them consistent and raises their accuracy to 0.738 (Caduceus-PH-1k) and 0.726 (augmented chunker), above MirrorDNA.

### 3.2 Accuracy at small scale

With 2.3M parameters, MirrorDNA has the highest average on Genomic Benchmarks (0.751) and on gene strand (0.956) (Table S17), but its lead over NT-v2 on Genomic Benchmarks is not significant (6 of 9 tasks, Wilcoxon *p* = 0.36; Table 3). On the NT tasks NT-v2 remains ahead (0.652 vs. 0.618, *p* = 0.001). MirrorDNA is not significantly different there from the 117M-parameter DNABERT-2 (0.631, *p* = 0.42), and more accurate than Caduceus-PS-1k and Caduceus-PH-1k, which had the same pretraining data and budget (0.595 and 0.579; *p* = 0.005 and 0.008). The centre readout matters most at splice sites, where the junction lies at the centre of the window. There MirrorDNA reaches 0.959–0.969, matching NT-v2 (0.959–0.963), against 0.792–0.856 with the pooled head. The remaining gap lies in two places. On the 12 histone and enhancer tasks, validation kept the pooled head, and NT-v2 and DNABERT-2 stay ahead. On Ensembl regulatory elements (0.758 vs. 0.901), all chunk-based models trail the single-nucleotide models. On gene strand, the orientation-aware head of MirrorDNA reaches 0.956, above every other model. Caduceus-PS-1k, used with its published invariant head, remains at chance (0.015), just as Proposition 1 predicts.

**Table 3.** Tasks won by MirrorDNA against each model (higher mean MCC over 3 seeds; not a per-task test), with the two-sided Wilcoxon signed-rank *p* over tasks. Gene strand has too few tasks for a test. All suites and SDs: Supplementary Tables S13 and S14.

| MirrorDNA vs. | NT (18) | GB (9) | Gene (2) | All (29) |
|---|---|---|---|---|
| NT-v2 | 3 (*p*=.001) | 6 (*p*=.36) | 1 | 10 (*p*=.076) |
| DNABERT-2 | 4 (*p*=.42) | 5 (*p*=.82) | 2 | 11 (*p*=.56) |
| HyenaDNA | 8 (*p*=.64) | 9 (*p*=.004) | 2 | 19 (*p*=.029) |
| Caduceus-PS | 14 (*p*=.005) | 7 (*p*=.20) | 2 | 23 (*p*<.001) |
| Caduceus-PH | 14 (*p*=.008) | 7 (*p*=.16) | 2 | 23 (*p*<.001) |
| Chunker + RC aug. | 8 (*p*=.93) | 6 (*p*=.30) | 2 | 16 (*p*=.21) |
| MirrorDNA-duplex | 12 (*p*=.018) | 6 (*p*=.30) | 2 | 20 (*p*=.002) |

### 3.3 Exact equivariance versus augmentation

MirrorDNA-duplex and the augmented chunker have the same architecture, pretraining data and budget. The only difference is whether the symmetry is built in or learned from augmented data (Fig. 4). Over 21 tasks (the 12 symmetric NT tasks and the 9 Genomic Benchmarks), both reach the same forward MCC when all training labels are used (0.595 vs. 0.587, *p* = 0.56). With 2,000 labels per task, the equivariant model is more accurate (0.479 vs. 0.462, a gain of 0.018), better on 16 of the 21 tasks (*p <* 0.001). With 500 labels the gain points the same way but is not significant ( +0.011, *p* = 0.15). At every budget the augmented model still changes its prediction with the strand for 10–12% of test windows. Its both-strands MCC is therefore 0.12–0.14 lower (all *p <* 10^−5^). Averaging the augmented model’s predictions over both strands removes this difference, but only at the output (0.379, 0.444 and 0.565 with 500, 2,000 and all labels, against 0.386, 0.453 and 0.563 for the equivariant model; all *p*≥ 0.16). The pretrained representations of non-equivariant models still differ between the two strands of the same DNA (Supplementary Table S15). We use the models frozen and look up the nearest neighbour of each reverse-complemented test sequence. For NT-v2, DNABERT-2, HyenaDNA, Caduceus-PH-1k and the augmented chunker, it is the same DNA for only 2–10% of test sequences. A linear probe trained on these embeddings changes 14–26% of its predictions with the strand. Its both-strands MCC is 0.148–0.273, against 0.449 for MirrorDNA and 0.441 for Caduceus-PS-1k, whose embeddings are identical on both strands. Augmentation during pretraining does not remove this: the augmented chunker flips 25.8% of the probe’s predictions. Frozen, MirrorDNA’s forward MCC (0.449) is within the range of the other models (0.436–0.476). Any use of these embeddings other than an averaged prediction, such as retrieval, clustering or a probe, therefore inherits the strand inconsistency.

**Fig. 4.**
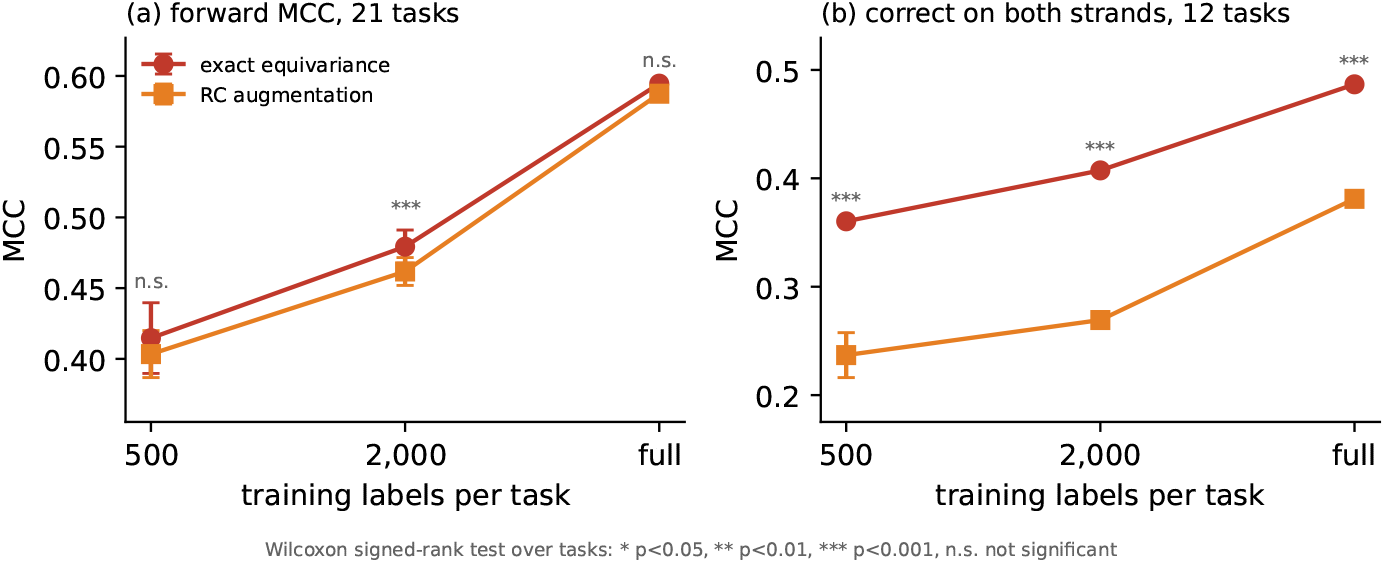
Exact equivariance versus augmentation at 500, 2,000 and all training labels per task (same architecture, pretraining data and budget). (a) Forward MCC over 21 tasks. (b) MCC counting only predictions correct on both strands, 12 strand-symmetric NT tasks. Mean ± SD over 3 seeds; Wilcoxon signed-rank test over tasks.

### 3.4 Longer inputs

MirrorDNA and Caduceus-PH-1k were both pretrained on 1-kb windows. On the gene-strand tasks with windows of up to 16 kb (Table 4), MirrorDNA loses little accuracy. From 1 to 16 kb, its MCC goes from 0.880 to 0.842 at transcription start sites and from 0.973 to 0.952 at splice donors. Caduceus-PH-1k is better at 1 kb on splice donors (0.988), but drops to 0.910 at 2 kb and 0.874 at 8 kb and 0.870 at 16 kb. At transcription start sites it falls from 0.728 to 0.645. At 8 kb, MirrorDNA is ahead by 0.20 MCC on start sites and 0.09 on donors. Because the informative signal sits near the centre of each window, this experiment measures robustness to input length rather than the use of distant context.

**Table 4.** Gene strand on longer inputs. (MCC, mean ± SD over 3 seeds). Both models were pretrained on 1-kb windows. Here MirrorDNA uses chunk features only (no base readout) at every length.

| Site | Model | 1 kb | 2 kb | 4 kb | 8 kb | 16 kb |
|---|---|---|---|---|---|---|
| TSS | Caduceus-PH-1k | 0.728<sub>±.038</sub> | 0.725<sub>±.034</sub> | 0.688<sub>±.029</sub> | 0.669<sub>±.016</sub> | 0.645<sub>±.017</sub> |
|  | MirrorDNA (ours) | **0.880**<sub>±.003</sub> | **0.872**<sub>±.012</sub> | **0.877**<sub>±.004</sub> | **0.871**<sub>±.004</sub> | **0.842**<sub>±.006</sub> |
| Donor | Caduceus-PH-1k | **0.988**<sub>±.002</sub> | 0.910<sub>±.010</sub> | 0.872<sub>±.003</sub> | 0.874<sub>±.003</sub> | 0.870<sub>±.004</sub> |
|  | MirrorDNA (ours) | 0.973<sub>±.003</sub> | **0.970**<sub>±.012</sub> | **0.968**<sub>±.004</sub> | **0.961**<sub>±.003</sub> | **0.952**<sub>±.004</sub> |

### 3.5 What the learned tokens capture

The tokenizer never sees an annotation, yet its boundaries follow gene structure (Fig. 5). MirrorDNA cuts about 1.7 times more often than background two bases before and two bases after splice junctions, and rarely at the junction itself. As a result, the chunk containing the first base of the intron at splice donors, read on the gene’s strand, is most often `AGGT`. This happens at 14.8% of sites, compared with 1.6% when the same cuts are shifted away from the site, a ninefold enrichment. The shorter `AGT` is enriched fifteenfold. `AGGT` spans the exon–intron junction and contains the invariant `GT` of the donor consensus, so the core of the donor signal ends up in a single token, on whichever strand the gene is read. Acceptors show a weaker version of the effect, and we found no consistent token at transcription start sites.

**Fig. 5.**
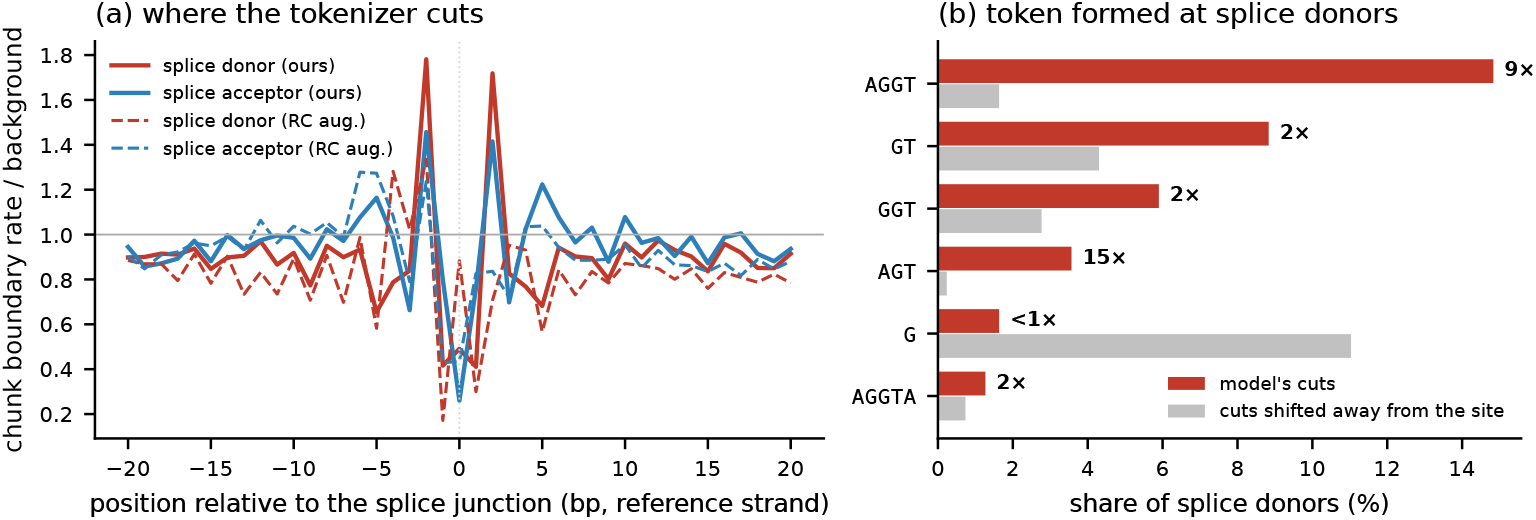
Tokens formed at splice sites. (held-out chromosomes 8 and 9). (a) Chunk-boundary rate around splice junctions relative to the background rate, for MirrorDNA and the augmented chunker. (b) Tokens containing the first intronic base at splice donors, read on the gene’s strand: share of sites with the model’s cuts and with the same cuts circularly shifted away from the site. Labels give the enrichment.

### 3.6 Case study: two genes on opposite strands

To see the strand behaviour on real genes, we looked at CDKN2A (chromosome 9, minus strand) and MYC (chro-mosome 8, plus strand). For each gene we ran the tokenizer on both strands and compared where it cuts. MirrorDNA cuts both strands at the same positions along the whole locus (Jaccard 1.000). The augmented chunker, which saw both strands during pretraining, does not: only 41% of its cut positions agree on CDKN2A (Fig. 6) and 35% on MYC (Supplementary Fig. S2). Around the first splice donor of each gene, MirrorDNA splits both strands into the same tokens, while the augmented chunker splits them differently (Fig. 6d).

**Fig. 6.**
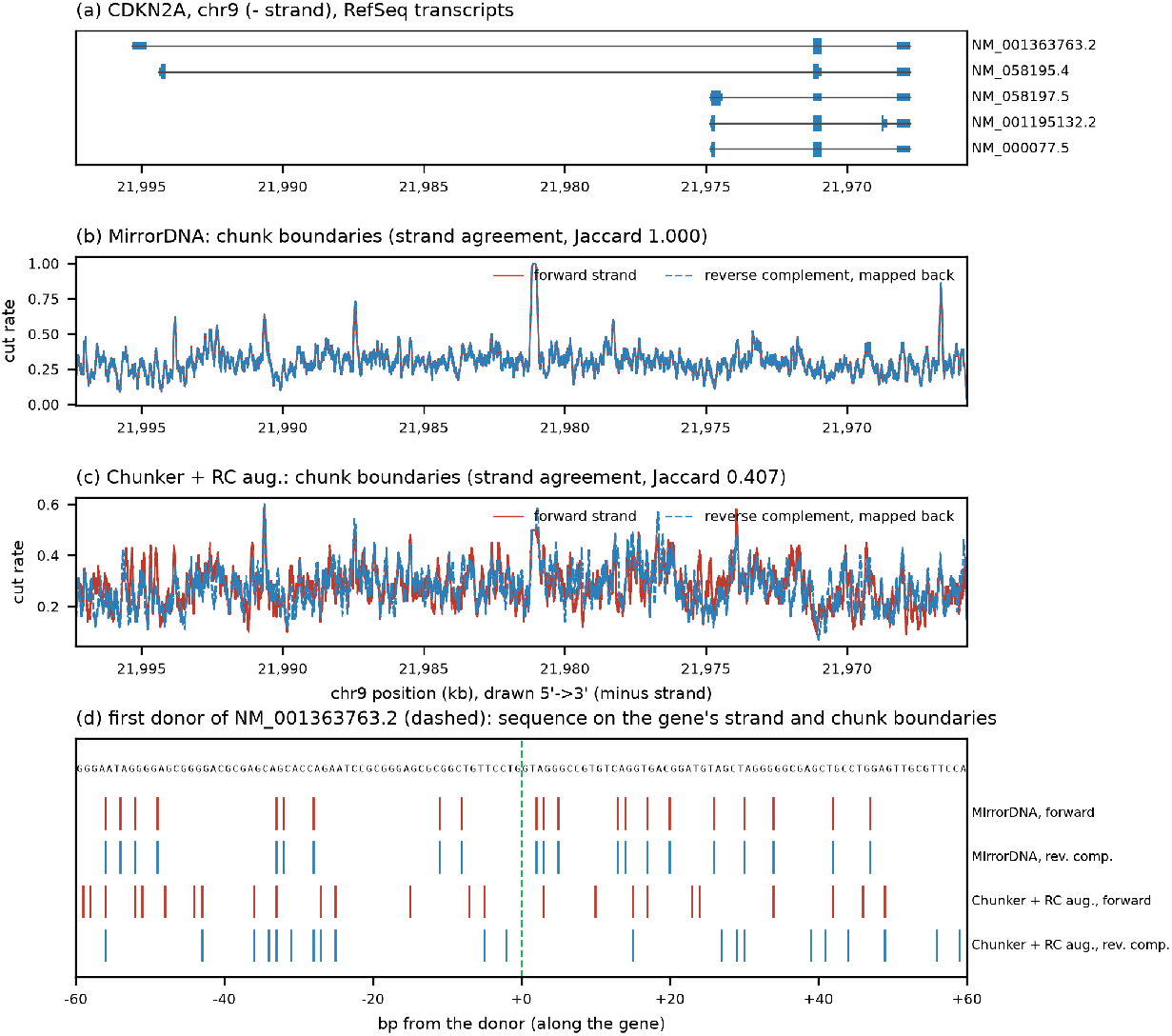
**CDKN2A** (chr9, minus strand; drawn 5′ →3 ′ along the gene). (a) RefSeq transcripts. (b, c) Chunk-boundary rate along the locus (100-bp moving average) from the forward strand and from the reverse complement mapped back, for MirrorDNA and for the augmented chunker; Jaccard = agreement of the cut positions between strands. (d) Sequence on the gene’s strand around the first donor of the longest transcript (dashed) and the cut positions on both strands, base by base.

## 4 Discussion

We have shown that learned tokenization and reverse-complement symmetry can be combined without approximation. This matters in practice, since widely used DNA language models change a quarter to a third of their predictions when the strand is flipped, and augmentation still leaves about one in ten. Both-strands accuracy and flip rates are easy to report, and we think they should become part of standard DNA benchmarks. Where Caduceus makes a single-nucleotide network equivariant, MirrorDNA makes the units of the sequence equivariant, and these units carry biological signal such as the splice-donor token.

### Limitations

MirrorDNA is small and was pretrained on human DNA only. It stays below NT-v2 on the NT tasks, mainly on histone marks, below the single-nucleotide models on Ensembl regulatory elements, and slightly below Caduceus on DeepSTARR. Scaling, multi-species data, variant effects and long-range tasks remain to be tested.

## 5 Conclusion

DNA has two strands, and for most regulatory labels it does not matter which one a model reads. Current DNA language models do not respect this. Their tokenizers cut the two strands of the same sequence differently, and their predictions change with the strand for a quarter to a third of test windows. We introduced MirrorDNA, whose learned tokenizer cuts both strands at mirrored positions, and proved that the whole model is equivariant to the reverse complement for any parameter values. Two heads on the same backbone handle both strand-symmetric and strand-dependent labels.

With 2.3M parameters and the pretraining budget of the 1-kb Caduceus models, MirrorDNA is more accurate than these models, and not significantly different from the 98M-parameter NT-v2 on Genomic Benchmarks or from the 117M-parameter DNABERT-2 on the Nucleotide Transformer tasks. It gives the same answer on both strands without test-time averaging, and keeps its accuracy on inputs sixteen times longer than in pretraining. Without any annotation, its tokenizer keeps the core splice-donor signal in a single token. More broadly, we hope that accuracy on both strands becomes a standard part of how DNA models are evaluated.

## Supplementary Material

Learning Strand-Equivariant Tokenizations of DNA with MirrorDNA Code, pretrained models, the per-seed results of every run and the scripts that produce every table and figure are available at https://github.com/NIRJHOR-DATTA/MirrorDNA

## S1 Strand-equivariance guarantees: statements and proofs

We show that the chunker’s outputs transform exactly with the reverse complement, for every parameter value. This holds at initialization, throughout training and without data augmentation. We then show that, for the risk on both strands and a convex loss, restricting to strand-invariant predictors does not exclude an optimal predictor. Group averaging (Lemma S4) is standard. The new ingredient is Lemma S2: the discrete, input-dependent segmentation itself commutes with the reverse complement.

*Notation*. Let Σ = {A, C, G, T} with complement *c* (A ↔ T, C ↔ G). The reverse complement of *x* ∈ Σ^*L*^ is ℛ (*x*)_*i*_ = *c*(*x*_*L*+1−*i*_), and ℛ ° ℛ = id. Hidden states are *H* ∈ ℝ^*L*×2*d*^, with each row split into two halves. Let *P* [*u*; *v*] = [*v*; *u*] swap the halves. The strand action on per-base states is *σ* (*H*)_*i*_ = *P H*_*L*+1−*i*_; on a sequence of *n* chunk vectors it is *σ*_*n*_ (*Z*)_*k*_ = *P Z*_*n*+1−*k*_ . Gap *j* ∈ {1, …, *L* − 1} lies between bases *j* and *j* + 1. The reversal of gap scores is (*J p*) _*j*_ = *p*_*L*− *j*_ . All three maps are involutions: 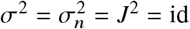.

### Definition S1

**(Equivariance)**. *A map F from sequences is* strand-equivariant *if F* (ℛ*x*) = *σF* (*x*), *and* strand-invariant *if F* (ℛ*x*) = *F* (*x*), *for all x*.

### Lemma S1

**(Encoder)**. *Let G map a pair of token sequences to a pair of per-base state sequences, G* (*x, x*^′^) = (*u, v*), *and suppose G commutes with exchanging its inputs: G* (*x*^′^, *x*) = (*v, u*). *Then the encoder H* (*x*) =*[u*; rev *v*] *with (u, v)* = *G (x, ℛx) is strand-equivariant. This covers the shared-weight convolutional encoder of our main model,H (x)* = [*g( x)* ; rev *g (ℛx) (no cross-strand terms), and the cross-strand gated duplex encoder used in MirrorDNA-duplex, provided its blocks update the two streams with the same weights*.

*Proof*. Let (*u, v*) = *G* (*x, ℛx*). Since *ℛ ℛx* = *x, G* (*ℛx, x*) = (*v, u*), so *H* (*ℛx*)_*i*_ = [ *v*_*i*_ ; *u*_*L*+1−*i*_ ] = *P* [ *u*_*L*+1−*i*_ ; *v*_*i*_ ] = *P H* (*x*)_*L*+1−*i*_ = *σ* (*H* (*x*))_*i*_. For the convolutional encoder, *G* (*x, x*^′^) = (*g*(*x*), *g*(*x*^′^)) commutes with the exchange. A duplex block maps (*u, v*) ↦ *F* (*u, v*), *F* (*v, u*) with one function *F* for both streams (each stream is gated by its own context and by the other stream’s context reversed into its coordinates), so exchanging *u* and *v* exchanges the two outputs. By induction over the blocks, *G* commutes with the exchange. □

### Lemma S2

(Mirrored segmentation).

*Let r* : ℝ^*L*×2*d*^→[ 0, 1] ^*L* −1^ *be any router, for example* 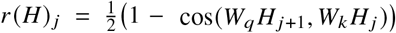, *with the cosine of a zero vector set to 0 (as an ϵ -stabilized normalization does), and define the symmetric router* 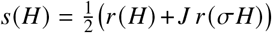. *Then s*(*σH*) = *J s*(*H*). *Consequently, with cuts* 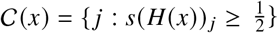,

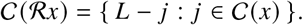

*The two strands are cut at exactly mirrored positions: they have the same number of chunks n, and chunk k of ℛx is chunk n* 1 *k of x, reverse-complemented*.

*Proof*. Using *σ*^2^ = id and *J*^2^ = id, 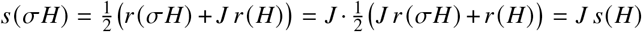. With Lemma S1,*s*(*H* (ℛ*x*))_*L*− *j*_ = *s*(*H* (*x*)) _*j*_, and the same threshold rule (including ties) is applied to equal scores. □

*Remark 1*. The raw router *r* is not mirror-symmetric, because *W*_*q*_ ≠ *W*_*k*_ and *r* compares a base with its left neighbour. Without the symmetrization in Lemma S2, the two strands are segmented differently and yield different tokens. Their representations are then not mirror images, and downstream predictions are not guaranteed to agree across strands. Consistency could still be imposed afterwards, for instance by averaging predictions over both strands, but this makes only the outputs consistent, not the representations (Section 3.3).

### Lemma S3

**(Chunk pooling and confidence)**. *Let Z* (*x*)_*k*_ *be the mean of H* (*x*)_*i*_ *over the bases i in chunk k, and let γ* (*x*)_*k*_ *be the mean of the router confidences v* _*j*_ = *s* _*j*_ *(for a cut) or* 1 − *s* _*j*_ *(no cut) over the gaps touching chunk k (inside it or on its boundary). Then Z* (ℛ*x*) = *σ*_*n*_ *Z* (*x*) *and γ* (ℛ*x*)_*n*+1−*k*_ = *γ* (*x*)_*k*_.

*Proof*. By Lemma S2, chunk *n*+ 1 −*k* of ℛ*x* contains exactly the bases *L+* 1− *i* for *i* in chunk *k* of *x*. By Lemma S1,*H( ℛ x)* _*L*_ *+*_1_− _*i*_ = *P H x* _*i*_, and a mean does not depend on the order of its terms, so *Z (ℛx)* _*n+* 1_− _*k*_ = *P Z (x)* _*k*_ . The gaps touching the two chunks are mirror images, with equal scores and equal cut decisions, so the confidences agree.

### Lemma S4

**(Group averaging)**. *For any map M on chunk (or base) sequences*, 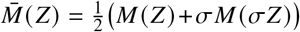 *satisfies*.

*Proof*. 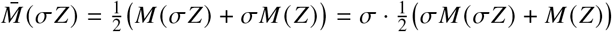. □

### Theorem S1

**(End-to-end strand equivariance)**. *Let the chunk transformer* 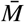 *and the decoder* 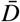 *be group-averaged as in Lemma S4, where the decoder map D* (*H, U*) *includes the residual projection of H, so that* 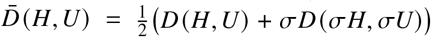. *Evaluation is deterministic (the model has no dropout). Let the upsampled states be* 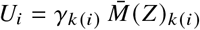, *where k* (*i*) *is the chunk containing base i. Let the language-model logits be* 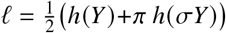, *where* 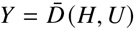, *h is any per-base output layer, and π reverses positions and complements the output letters. Then,for every parameter value and every input x:*

1. *the per-base predictions are strand-equivariant: l*(ℛ*x*) = *π l*(*x*);
2. *the sequence features ϕ*(*x*) = [*a* + *b*, |*a* − *b*|], *where* [*a*; *b*] *is the mean (or max) of* 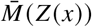 *over chunks, are strand-invariant: ϕ (ℛ)x* = *ϕ (x), and so are the same combinations of the mean and max of Y over all bases (base readout) or over any set of positions that is mapped onto itself by i* ↦*L +*1 −*i, such as the central window of the input (centre readout);*
3. *any classifier or regressor f* = *q* ° *ϕ satisfies f* (*ℛx*) = *f* (*x*), *so in exact arithmetic its flip rate is zero on any test set;*
4. *the orientation-aware features are exchanged:* [*a*; *b*] (*ℛx*) = [*b*; *a*] (*x*).

*Proof*. By Lemmas S3 and S4, 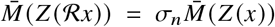. Combined with the mirrored chunk assignment and confidences, this gives *U* (ℛ*x*) = *σU* (*x*). Lemma S4 applied to 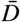, with *H* and *U* transforming together, gives *Y* (ℛ*x*) = *σY* (*x*). For the output layer, 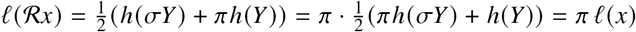, since *π*^2^ = id.This proves (1). For (2), pooling over chunks (or over bases, or over a set of positions mapped onto itself by *i* ↦→ *L* +1−*i*, using *Y* (R*x*) = *σY* (*x*)) ignores their order, and *σ*_*n*_ swaps the halves, *a* ↔ *b*. Both *a* + *b* and |*a* − *b*| are symmetric in (*a, b*). (3) follows from (2), and (4) is the exchange *a* ↔ *b* itself. □

*Remark 2 (Pretraining and numerics)*. During masked-language-model pretraining, cuts are forced around masked bases and the corresponding gaps get confidence 1. The identities then apply to the corrupted input itself: if 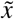 is the corrupted version of *x*, part (1) relates 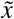 and 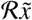, which has the mask positions mirrored and the randomly replaced bases complemented, with the forced cuts mirrored accordingly. The identities are exact in real arithmetic. In floating point the two strands agree up to rounding, with a relative error of about 10^−7^ in 32-bit precision, so a prediction that lies within rounding error of the decision threshold can still flip. Evaluated in 32-bit precision, this happens for 0.003% of the strand-symmetric test windows of MirrorDNA.

### Invariance and the risk on both strands

Let *D* be the test distribution over pairs (*x, y)*, and let ℛ*D* be the distribution of ( ℛ*x, y)* . For tasks whose label does not depend on the strand read (histone marks, enhancers), a sequence and its reverse complement are equally valid test inputs. The relevant objective is then the *strand-robust risk*

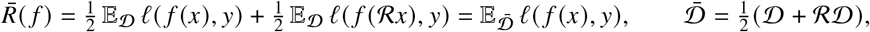

whose empirical counterpart is the average of the per-example losses on the forward and the reverse-complemented test set. This is a statement about convex losses, not about MCC or our both-strands scoring rule, which are nonlinear functions of the confusion matrix. Standard benchmarks estimate only the first term. When *D*≠ℛ *D*, for example when informative content sits off-centre in the benchmark windows, that term can reward orientation-dependent shortcuts that do not transfer to the other strand.

#### Proposition S1

**(Invariance is free for the strand-robust risk)**. *Let predictions take values in a convex set (class probabilities or real numbers), and let l be convex in its first argument (for example, cross-entropy on predicted probabilities, or squared error)*.

1. *For any predictor f, its strand average*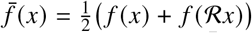 *is strand-invariant and satisfies* 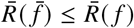.
2. *There exists a strand-invariant Bayes-optimal predictor for* 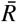, *since* 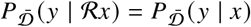.

*Hence restricting to strand-invariant predictors does not exclude an optimal predictor for* 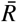, *and any predictor can be matched or improved by its strand average. The result concerns unrestricted predictors, or hypothesis classes closed under strand averaging. It does not imply that the particular invariant features of Theorem S1 lose no information. For instance, [a +b*,| *a* − *b\* ]*maps (a, b)* = ((0, 0 ), (1, 1)) *and (a, b )*= ((0, 1),( 1, 0)) *to the same features, although the two are not related by a strand swap*.

*Proof*. (1) Invariance holds because ℛ ℛ *x* = *x*. By convexity (Jensen’s inequality),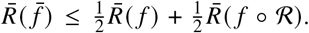 Moreover 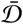 is invariant under (*x, y*) ↦ (ℛ *x, y*), so 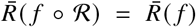 The same invariance of 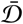 gives 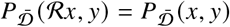 for all (*x, y*). The conditional of *y* is therefore equal at *x* and ℛ *x*. □

*Remark 3 (Scope)*. Proposition S1 assumes the label is strand-symmetric. For direction-dependent labels (promoters, splice donors and acceptors), the correct symmetry is equivariance with a relabelling rather than invariance. Our backbone lets a head learn such labels by keeping the orientation-aware features [*a*; *b*] instead of *ϕ* (main text, Section 2.3), which are swapped by the reverse complement. We do not constrain the head to output the opposite label on the other strand.

### S2 Implementation and training details

#### Router training

As in H-Net, each chunk carries a confidence, the mean of *s* _*j*_ over the cut gaps and of 1 −*s* _*j*_ over the uncut gaps that touch it. It is applied with a straight-through estimator (forward value 1), so the router receives a gradient. Mirrored chunks touch mirrored gaps with equal scores and equal decisions, so their confidences are equal and the guarantee is kept. The H-Net ratio loss (weight 0.03) keeps the fraction of cut gaps near 1 /*R* for a target chunk length of *R* = 4 bp. The cosine in the router is computed on *ϵ* -normalized vectors, so it is defined for every input.

#### Pretraining

Masking follows BERT. Of the 15% selected bases, 80% are replaced by a mask token, 10% by a random base and 10% are left unchanged. Masked bases are always chunk boundaries on both sides. The guarantees apply to the corrupted input itself, as explained after Theorem S1. Soft-masked repeats are weighted 0.1 in the loss. We use AdamW with *β* = (0.9, 0.95), weight decay 0.1 and a peak learning rate of 1.5 ×10^−3^, with 1,000 warm-up steps, cosine decay to 10% of the peak, gradient clipping at 1.0 and bfloat16 autocast. The main model has *d* = 96, four encoder blocks, four transformer layers with four heads and two decoder blocks, for 2.35M parameters. Its target chunk length is 4 bp, and chunks average 3.1 bp after training. MirrorDNA-duplex is the same model with a cross-strand gated encoder (2.50M parameters). The augmented chunker has the architecture of MirrorDNA-duplex without symmetrization (2.52M parameters) and was pretrained with reverse-complement augmentation, on the same data and budget. No model uses dropout.

#### Fine-tuning

We use AdamW (weight decay 0.01) with a linear warm-up over the first 5% of steps, cosine decay to 10% of the peak and gradient clipping at 1.0. The batch size is 64 for MirrorDNA and 32 for the baselines, whose long inputs are split into micro-batches with gradient accumulation. Training runs for at most 10 epochs with early stopping on validation MCC (patience 3), and the best epoch is evaluated on the test set. We use seeds 0, 1 and 2, and a random 10% of each official training set for validation. On direction-dependent tasks the augmented models are fine-tuned without augmentation, since flipping the strand while keeping the label would teach them to ignore it. In the main results MirrorDNA pools both the chunk features and the per-base states *Y* (base readout), and for each task the pooled head or the centre readout is chosen by mean validation MCC over the three seeds (Section 2.4). The length experiment uses chunk features only.

#### Baseline

NT-v2 is `InstaDeepAI/nucleotide-transformer-v2-100m-multi-species` with 6-mer tokens and learning rate 3 × 10^−5^. DNABERT-2 is `zhihan1996/DNABERT-2-117M` with learning rate 3 × 10^−5^, run with `transformers` 4.46.3 and its Triton attention kernel disabled. HyenaDNA is `LongSafari/hyenadna-small-32k -seqlen-hf`, run in full precision. Caduceus-PS-1k and Caduceus-PH-1k are `yairschiff/caduceus-ps_seqlen-1k_d_model-118_n_layer-4_lr-8e-3` and its `ph` counterpart. They were pretrained with the same data and token budget as MirrorDNA. We load them with the model code of the 131k checkpoints, which has the same architecture and parameter names, and stop the run if any weight is missing. Caduceus-PS uses its published invariant head, which averages the pooled features of the two strands. Caduceus-PH follows its published protocol, with reverse-complement augmentation on the NT and Genomic Benchmarks tasks, and is fine-tuned without augmentation on the gene-strand tasks. Every baseline uses the same mean-and-max pooled linear head.

#### Gene-strand tasks

Windows are centred on the transcription start sites and splice donors of RefSeq protein-coding (NM_) transcripts of hg38 and labelled with the strand of the transcript. Chromosomes 8 and 9 are held out for testing. The same sites are used at 1, 2, 4, 8 and 16 kb.

#### DeepSTAR

We use the official training, validation and test sets of 249-bp sequences (402,296, 40,570 and 41,186 sequences). Targets are standardized with the training mean and standard deviation. The loss is the mean-squared error over the two targets, and the model is selected on the mean validation Pearson’s *r*. All other settings are as above. The augmented models (Caduceus-PH-1k and the augmented chunker) are fine-tuned with reverse-complement augmentation, which is valid here because enhancer activity does not depend on the strand read.

#### Numerical precision

The identities of Theorem S1 are exact in real arithmetic. In 32-bit precision the two strands agree to a relative error of about 10^−7^, so a prediction that lies within rounding error of the decision threshold can occasionally flip. Evaluated in 32-bit precision, MirrorDNA changes its prediction for 0.003% of the strand-symmetric test windows, and MirrorDNA-duplex for 0.004%.

### S3 Detailed results

#### S3.1 Accuracy per task

Forward MCC on the official test sets, mean ±SD over three seeds (one fine-tuning protocol for all models; for MirrorDNA the head is chosen per task on validation, Section 2.4). Model names: Cad.-PS/PH = Caduceus-PS/PH-1k; Chunk.+aug. = learned chunker with reverse-complement augmentation; MDNA-dup. = MirrorDNA-duplex; HyenaDNA = HyenaDNA-small.

### S3.2 Accuracy on both strands, per task

**Table S1.** Nucleotide Transformer benchmark (revised), 18 tasks: forward MCC. MirrorDNA: head chosen per task on validation;^*c*^ centre readout chosen (Section 2.4).

| Task | NT-v2 | DNABERT-2 | HyenaDNA | HyenaDNA-1k | Cad.-PS | Cad.-PH | Chunk.+aug. | MDNA-dup. | MirrorDNA |
|---|---|---|---|---|---|---|---|---|---|
| H2AFZ | .495<sub>±.026</sub> | .521<sub>±.010</sub> | .490<sub>±.007</sub> | .492<sub>±.015</sub> | .426<sub>±.001</sub> | .422<sub>±.032</sub> | .446<sub>±.012</sub> | .425<sub>±.005</sub> | .422<sub>±.016</sub> |
| H3K27ac | .502<sub>±.008</sub> | .510<sub>±.020</sub> | .464<sub>±.009</sub> | .463<sub>±.012</sub> | .435<sub>±.018</sub> | .444<sub>±.006</sub> | .465<sub>±.030</sub> | .443<sub>±.003</sub> | .446<sub>±.017</sub> |
| H3K27me3 | .585<sub>±.007</sub> | .607<sub>±.005</sub> | .574<sub>±.007</sub> | .568<sub>±.005</sub> | .546<sub>±.005</sub> | .545<sub>±.001</sub> | .544<sub>±.006</sub> | .541<sub>±.010</sub> | .541<sub>±.011</sub> |
| H3K36me3 | .618<sub>±.011</sub> | .632<sub>±.002</sub> | .563<sub>±.010</sub> | .566<sub>±.004</sub> | .531<sub>±.005</sub> | .538<sub>±.004</sub> | .537<sub>±.008</sub> | .532<sub>±.015</sub> | .543<sub>±.010</sub> |
| H3K4me1 | .491<sub>±.002</sub> | .501<sub>±.004</sub> | .453<sub>±.011</sub> | .460<sub>±.002</sub> | .414<sub>±.013</sub> | .416<sub>±.019</sub> | .425<sub>±.019</sub> | .432<sub>±.010</sub> | .423<sub>±.013</sub> |
| H3K4me2 | .557<sub>±.007</sub> | .575<sub>±.009</sub> | .519<sub>±.008</sub> | .537<sub>±.009</sub> | .469<sub>±.013</sub> | .479<sub>±.006</sub> | .496<sub>±.017</sub> | .441<sub>±.015</sub> | .484<sub>±.013</sub> |
| H3K4me3 | .613<sub>±.011</sub> | .615<sub>±.014</sub> | .626<sub>±.006</sub> | .600<sub>±.017</sub> | .596<sub>±.005</sub> | .608<sub>±.008</sub> | .609<sub>±.010</sub> | .601<sub>±.005</sub> | .600<sub>±.011</sub> |
| H3K9ac | .526<sub>±.005</sub> | .544<sub>±.022</sub> | .516<sub>±.009</sub> | .535<sub>±.027</sub> | .518<sub>±.023</sub> | .519<sub>±.001</sub> | .523<sub>±.013</sub> | .495<sub>±.034</sub> | .524<sub>±.012</sub> |
| H3K9me3 | .445<sub>±.019</sub> | .469<sub>±.020</sub> | .416<sub>±.008</sub> | .422<sub>±.015</sub> | .394<sub>±.024</sub> | .382<sub>±.012</sub> | .398<sub>±.029</sub> | .411<sub>±.026</sub> | .375<sub>±.039</sub> |
| H4K20me1 | .637<sub>±.011</sub> | .440<sub>±.381</sub> | .628<sub>±.003</sub> | .619<sub>±.015</sub> | .580<sub>±.008</sub> | .580<sub>±.010</sub> | .578<sub>±.013</sub> | .579<sub>±.008</sub> | .591<sub>±.008</sub> |
| enhancers | .524<sub>±.004</sub> | .533<sub>±.008</sub> | .496<sub>±.005</sub> | .496<sub>±.010</sub> | .492<sub>±.010</sub> | .485<sub>±.009</sub> | .494<sub>±.011</sub> | .488<sub>±.002</sub> | .492<sub>±.017</sub> |
| enhancers_types | .479<sub>±.008</sub> | .502<sub>±.013</sub> | .431<sub>±.013</sub> | .459<sub>±.008</sub> | .442<sub>±.017</sub> | .447<sub>±.003</sub> | .463<sub>±.004</sub> | .452<sub>±.016</sub> | .450<sub>±.014</sub> |
| promoter_all | .746<sub>±.010</sub> | .748<sub>±.001</sub> | .720<sub>±.008</sub> | .723<sub>±.014</sub> | .719<sub>±.004</sub> | .707<sub>±.006</sub> | .735<sub>±.004</sub> | .732<sub>±.010</sub> | .743<sub>±.009</sub> |
| promoter_no_tata | .757<sub>±.010</sub> | .769<sub>±.007</sub> | .737<sub>±.006</sub> | .750<sub>±.014</sub> | .737<sub>±.008</sub> | .732<sub>±.005</sub> | .750<sub>±.015</sub> | .731<sub>±.008</sub> | .765<sub>±.006</sub> |
| promoter_tata | .875<sub>±.005</sub> | .846<sub>±.024</sub> | .833<sub>±.015</sub> | .825<sub>±.011</sub> | .834<sub>±.044</sub> | .682<sub>±.010</sub> | .837<sub>±.006</sub> | .816<sub>±.033</sub> | .835<sub>±.024</sub> |
| splice_sites_acceptors | .961<sub>±.002</sub> | .835<sub>±.008</sub> | .785<sub>±.087</sub> | .848<sub>±.011</sub> | .842<sub>±.007</sub> | .829<sub>±.018</sub> | .726<sub>±.019</sub> | .739<sub>±.027</sub> | .959<sup>*c*</sup><sub>±.004</sub> |
| splice_sites_all | .959<sub>±.005</sub> | .859<sub>±.007</sub> | .900<sub>±.009</sub> | .884<sub>±.004</sub> | .900<sub>±.004</sub> | .795<sub>±.042</sub> | .766<sub>±.007</sub> | .825<sub>±.026</sub> | .963<sup>*c*</sup><sub>±.002</sub> |
| splice_sites_donors | .963<sub>±.001</sub> | .857<sub>±.004</sub> | .909<sub>±.028</sub> | .923<sub>±.016</sub> | .836<sub>±.046</sub> | .818<sub>±.029</sub> | .773<sub>±.009</sub> | .812<sub>±.002</sub> | .969<sup>*c*</sup><sub>±.004</sub> |

**Table S2.** Genomic Benchmarks, 9 tasks: forward MCC. MirrorDNA: head chosen per task on validation; ^*c*^ centre readout chosen (Section 2.4).

| Task | NT-v2 | DNABERT-2 | HyenaDNA | HyenaDNA-1k | Cad.-PS | Cad.-PH | Chunk.+aug. | MDNA-dup. | MirrorDNA |
|---|---|---|---|---|---|---|---|---|---|
| coding vs intergenomic | .891<sub>±.004</sub> | .896<sub>±.005</sub> | .821<sub>±.003</sub> | .820<sub>±.004</sub> | .825<sub>±.004</sub> | .813<sub>±.002</sub> | .881<sub>±.001</sub> | .884<sub>±.002</sub> | .882<sub>±.000</sub> |
| human or worm | .936<sub>±.001</sub> | .952<sub>±.002</sub> | .926<sub>±.002</sub> | .925<sub>±.001</sub> | .940<sub>±.002</sub> | .940<sub>±.002</sub> | .941<sub>±.002</sub> | .940<sub>±.001</sub> | .944<sub>±.001</sub> |
| Drosophila enhancers | .591<sub>±.012</sub> | .326<sub>±.272</sub> | .462<sub>±.037</sub> | .134<sub>±.191</sub> | .499<sub>±.075</sub> | .531<sub>±.034</sub> | .622<sub>±.085</sub> | .629<sub>±.014</sub> | .628<sub>±.016</sub> |
| mouse enhancers | .675<sub>±.044</sub> | .440<sub>±.383</sub> | .626<sub>±.005</sub> | .603<sub>±.029</sub> | .614<sub>±.009</sub> | .563<sub>±.024</sub> | .533<sub>±.092</sub> | .680<sub>±.018</sub> | .708<sup>*c*</sup><sub>±.033</sub> |
| enhancers (Cohn) | .482<sub>±.009</sub> | .520<sub>±.004</sub> | .418<sub>±.002</sub> | .448<sub>±.012</sub> | .490<sub>±.003</sub> | .492<sub>±.010</sub> | .488<sub>±.008</sub> | .472<sub>±.006</sub> | .473<sub>±.002</sub> |
| enhancers (Ensembl) | .822<sub>±.007</sub> | .832<sub>±.014</sub> | .742<sub>±.004</sub> | .714<sub>±.002</sub> | .825<sub>±.005</sub> | .775<sub>±.002</sub> | .780<sub>±.003</sub> | .830<sub>±.004</sub> | .838<sub>±.005</sub> |
| Ensembl regulatory | .901<sub>±.004</sub> | .819<sub>±.006</sub> | .619<sub>±.205</sub> | .821<sub>±.004</sub> | .811<sub>±.036</sub> | .831<sub>±.007</sub> | .577<sub>±.038</sub> | .677<sub>±.019</sub> | .758<sup>*c*</sup><sub>±.011</sub> |
| non-TATA promoters | .812<sub>±.008</sub> | .893<sub>±.008</sub> | .822<sub>±.009</sub> | .883<sub>±.007</sub> | .897<sub>±.012</sub> | .852<sub>±.007</sub> | .910<sub>±.004</sub> | .908<sub>±.001</sub> | .902<sub>±.005</sub> |
| open chromatin | .562<sub>±.007</sub> | .613<sub>±.007</sub> | .516<sub>±.013</sub> | .536<sub>±.005</sub> | .621<sub>±.003</sub> | .618<sub>±.005</sub> | .626<sub>±.002</sub> | .624<sub>±.005</sub> | .624<sub>±.007</sub> |

**Table S3.**
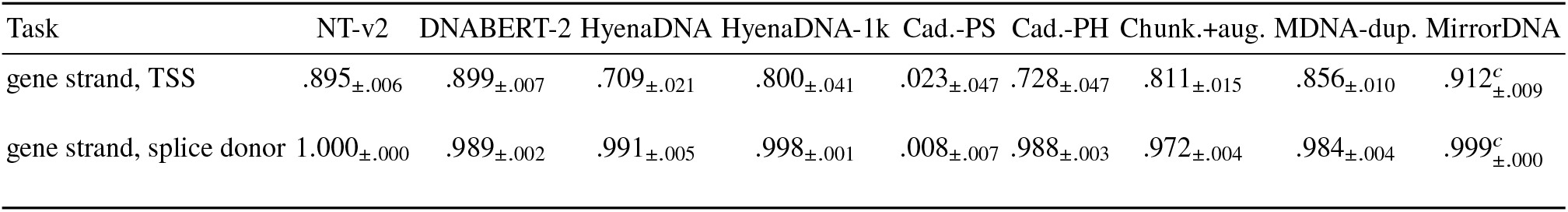
Gene-strand tasks (1-kb windows): forward MCC. MirrorDNA: head chosen per task on validation; ^*c*^ centre readout chosen (Section 2.4).

| Task | NT-v2 | DNABERT-2 | HyenaDNA | HyenaDNA-1k | Cad.-PS | Cad.-PH | Chunk.+aug. | MDNA-dup. | MirrorDNA |
|---|---|---|---|---|---|---|---|---|---|
| gene strand, TSS | .895<sub>±.006</sub> | .899<sub>±.007</sub> | .709<sub>±.021</sub> | .800<sub>±.041</sub> | .023<sub>±.047</sub> | .728<sub>±.047</sub> | .811<sub>±.015</sub> | .856<sub>±.010</sub> | .912<sup>*c*</sup><sub>±.009</sub> |
| gene strand, splice donor | 1.000<sub>±.000</sub> | .989<sub>±.002</sub> | .991<sub>±.005</sub> | .998<sub>±.001</sub> | .008<sub>±.007</sub> | .988<sub>±.003</sub> | .972<sub>±.004</sub> | .984<sub>±.004</sub> | .999<sup>*c*</sup><sub>±.000</sub> |

**Table S4.** Suite averages (forward MCC; SD of the per-seed suite average). MirrorDNA: head chosen per task on validation.

| Model | NT-18 | NT 12 sym. | NT 6 dir. | GB-9 | Gene strand |
|---|---|---|---|---|---|
| NT-v2 | .652<sub>±.002</sub> | .539<sub>±.004</sub> | .877<sub>±.005</sub> | .741<sub>±.004</sub> | .947<sub>±.003</sub> |
| DNABERT-2 | .631<sub>±.018</sub> | .537<sub>±.030</sub> | .819<sub>±.006</sub> | .699<sub>±.038</sub> | .944<sub>±.003</sub> |
| HyenaDNA-small | .615<sub>±.007</sub> | .515<sub>±.001</sub> | .814<sub>±.018</sub> | .661<sub>±.021</sub> | .850<sub>±.013</sub> |
| HyenaDNA-tiny-1k | .621<sub>±.001</sub> | .518<sub>±.002</sub> | .826<sub>±.004</sub> | .654<sub>±.020</sub> | .899<sub>±.020</sub> |
| Caduceus-PS-1k | .595<sub>±.002</sub> | .487<sub>±.002</sub> | .812<sub>±.007</sub> | .725<sub>±.003</sub> | .015<sub>±.027</sub> |
| Caduceus-PH-1k | .579<sub>±.003</sub> | .489<sub>±.004</sub> | .761<sub>±.009</sub> | .713<sub>±.005</sub> | .858<sub>±.023</sub> |
| Chunker + RC aug. | .587<sub>±.005</sub> | .498<sub>±.006</sub> | .765<sub>±.004</sub> | .706<sub>±.005</sub> | .891<sub>±.006</sub> |
| MirrorDNA duplex | .583<sub>±.002</sub> | .487<sub>±.002</sub> | .776<sub>±.009</sub> | .738<sub>±.003</sub> | .920<sub>±.004</sub> |
| MirrorDNA (ours) | .618<sub>±.003</sub> | .491<sub>±.004</sub> | .872<sub>±.006</sub> | .751<sub>±.002</sub> | .956<sub>±.004</sub> |

**Table S5.** MirrorDNA against every model on the 12 strand-symmetric NT tasks, forward and on both strands: mean MCC difference over tasks (MirrorDNA minus baseline), tasks won/lost, two-sided Wilcoxon signed-rank test over tasks, and tasks with a significant per-task Welch test over seeds after Holm correction (won/lost). On these tasks validation kept MirrorDNA’s pooled head. Tests on all suites: Supplementary Table S14.

| Suite | Baseline | Metric | Δ | W/L | *p* | sig. W/L |
|---|---|---|---|---|---|---|
| NT 12 sym. | NT-v2 | forward | -0.049 | 0/12 | 0.00049 | 0/3 |
| NT 12 sym. | DNABERT-2 | forward | -0.047 | 1/11 | 0.034 | 0/5 |
| NT 12 sym. | HyenaDNA-small | forward | -0.024 | 2/10 | 0.0068 | 0/0 |
| NT 12 sym. | HyenaDNA-tiny-1k | forward | -0.027 | 0/12 | 0.00049 | 0/0 |
| NT 12 sym. | Caduceus-PS-1k | forward | +0.004 | 8/4 | 0.15 | 0/0 |
| NT 12 sym. | Caduceus-PH-1k | forward | +0.002 | 8/4 | 0.34 | 0/0 |
| NT 12 sym. | Chunker + RC aug. | forward | -0.007 | 3/9 | 0.064 | 0/0 |
| NT 12 sym. | MirrorDNA duplex | forward | +0.004 | 6/6 | 0.47 | 0/0 |
| NT 12 sym. | NT-v2 | both strands | +0.338 | 12/0 | 0.00049 | 12/0 |
| NT 12 sym. | DNABERT-2 | both strands | +0.303 | 12/0 | 0.00049 | 7/0 |
| NT 12 sym. | HyenaDNA-small | both strands | +0.411 | 12/0 | 0.00049 | 12/0 |
| NT 12 sym. | HyenaDNA-tiny-1k | both strands | +0.410 | 12/0 | 0.00049 | 12/0 |
| NT 12 sym. | Caduceus-PS-1k | both strands | +0.004 | 8/4 | 0.15 | 0/0 |
| NT 12 sym. | Caduceus-PH-1k | both strands | +0.102 | 12/0 | 0.00049 | 12/0 |
| NT 12 sym. | Chunker + RC aug. | both strands | +0.110 | 12/0 | 0.00049 | 12/0 |
| NT 12 sym. | MirrorDNA duplex | both strands | +0.004 | 6/6 | 0.47 | 0/0 |

### S3.3 Longer inputs

**Table S6.** Gene strand at increasing window length (MCC, mean ±SD over seeds; *n* = seeds). The length experiment uses chunk features only. Rows marked +base use the pooled head with base readout; in Table S17 validation chose the centre readout for both gene-strand tasks (0.912 and 0.999).

| model | site | 1024 bp | 2048 bp | 4096 bp | 8192 bp | 16384 bp |
|---|---|---|---|---|---|---|
| MirrorDNA (ours) | TSS | 0.880 +- 0.003 (n=3) | 0.872 +- 0.012 (n=3) | 0.877 +- 0.004 (n=3) | 0.871 +- 0.004 (n=3) | 0.842 +- 0.006 (n=3) |
| MirrorDNA (ours) | donor | 0.973 +- 0.003 (n=3) | 0.970 +- 0.012 (n=3) | 0.968 +- 0.004 (n=3) | 0.961 +- 0.003 (n=3) | 0.952 +- 0.004 (n=3) |
| MirrorDNA (ours) +base | TSS | 0.857 +- 0.016 (n=3) | - | - | - | - |
| MirrorDNA (ours) +base | donor | 0.987 +- 0.003 (n=3) | - | - | - | - |
| Caduceus-PH-1k | TSS | 0.728 +- 0.038 (n=3) | 0.725 +- 0.034 (n=3) | 0.688 +- 0.029 (n=3) | 0.669 +- 0.016 (n=3) | 0.645 +- 0.017 (n=3) |
| Caduceus-PH-1k | donor | 0.988 +- 0.002 (n=3) | 0.910 +- 0.010 (n=3) | 0.872 +- 0.003 (n=3) | 0.874 +- 0.003 (n=3) | 0.870 +- 0.004 (n=3) |
| Caduceus-PS-1k | TSS | 0.023 +- 0.039 (n=3) | - | - | - | - |
| Caduceus-PS-1k | donor | 0.008 +- 0.006 (n=3) | - | - | - | - |

### S3.4 Regression: DeepSTARR

MirrorDNA uses the pooled head (no centre readout), since enhancer activity is a property of the whole sequence.

**Table S7.**
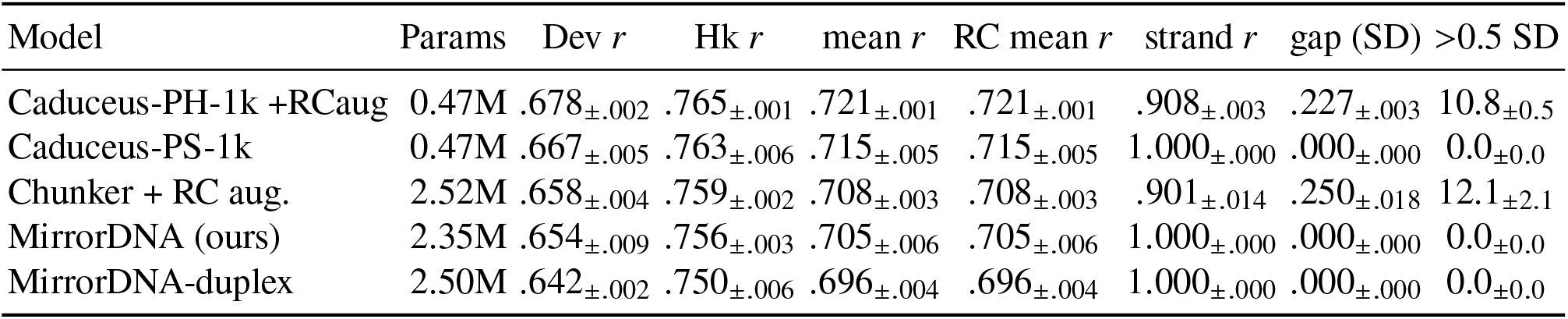
DeepSTARR enhancer-activity regression (official test set; mean ±SD over 3 seeds). Pearson’s *r* for developmental (Dev) and housekeeping (Hk) activity, their mean on the test set as given and reverse-complemented (RC); strand *r*: correlation of the predictions for a sequence and its reverse complement; gap: mean absolute difference of the two predictions in target SD units; *>*0.5 SD: % of sequences whose two predictions differ by more than half an SD. Published DeepSTARR network: Dev 0.68, Hk 0.74.

| Model | Params | Dev *r* | Hk *r* | mean *r* | RC mean *r* | strand *r* | gap (SD) | >0.5 SD |
|---|---|---|---|---|---|---|---|---|
| Caduceus-PH-1k +RCaug | 0.47M | .678<sub>±.002</sub> | .765<sub>±.001</sub> | .721<sub>±.001</sub> | .721<sub>±.001</sub> | .908<sub>±.003</sub> | .227<sub>±.003</sub> | 10.8<sub>±0.5</sub> |
| Caduceus-PS-1k | 0.47M | .667<sub>±.005</sub> | .763<sub>±.006</sub> | .715<sub>±.005</sub> | .715<sub>±.005</sub> | 1.000<sub>±.000</sub> | .000<sub>±.000</sub> | 0.0<sub>±0.0</sub> |
| Chunker + RC aug. | 2.52M | .658<sub>±.004</sub> | .759<sub>±.002</sub> | .708<sub>±.003</sub> | .708<sub>±.003</sub> | .901<sub>±.014</sub> | .250<sub>±.018</sub> | 12.1<sub>±2.1</sub> |
| MirrorDNA (ours) | 2.35M | .654<sub>±.009</sub> | .756<sub>±.003</sub> | .705<sub>±.006</sub> | .705<sub>±.006</sub> | 1.000<sub>±.000</sub> | .000<sub>±.000</sub> | 0.0<sub>±0.0</sub> |
| MirrorDNA-duplex | 2.50M | .642<sub>±.002</sub> | .750<sub>±.006</sub> | .696<sub>±.004</sub> | .696<sub>±.004</sub> | 1.000<sub>±.000</sub> | .000<sub>±.000</sub> | 0.0<sub>±0.0</sub> |

### S3.5 Tokenization of the two strands

**Table S8.** Tokenization of JASPAR 2024 motif occurrences (chromosomes 8 and 9): mean token length, tokens per motif occurrence, the same relative to random cuts of equal density, % of occurrences kept in one token, % tokenized differently on the two strands, and the mean Jaccard index of the two strands’ cut sets in 2,000 random 1-kb windows.

| Tokenizer | bp/token | tokens/motif | vs. random | whole (%) | differ (%) | Jaccard |
|---|---|---|---|---|---|---|
| MirrorDNA (ours) | 3.19 | 3.72 | 0.99 | 8.8 | 0.0 | 1.000 |
| Chunker + RC aug. | 3.57 | 3.40 | 0.98 | 11.5 | 87.8 | 0.393 |
| BPE (DNABERT-2) | 5.15 | 2.73 | 1.00 | 3.4 | 92.1 | – |
| 6-mer (NT) | 5.98 | 2.48 | 1.00 | 1.9 | 100.0 | – |

**Table S9.**
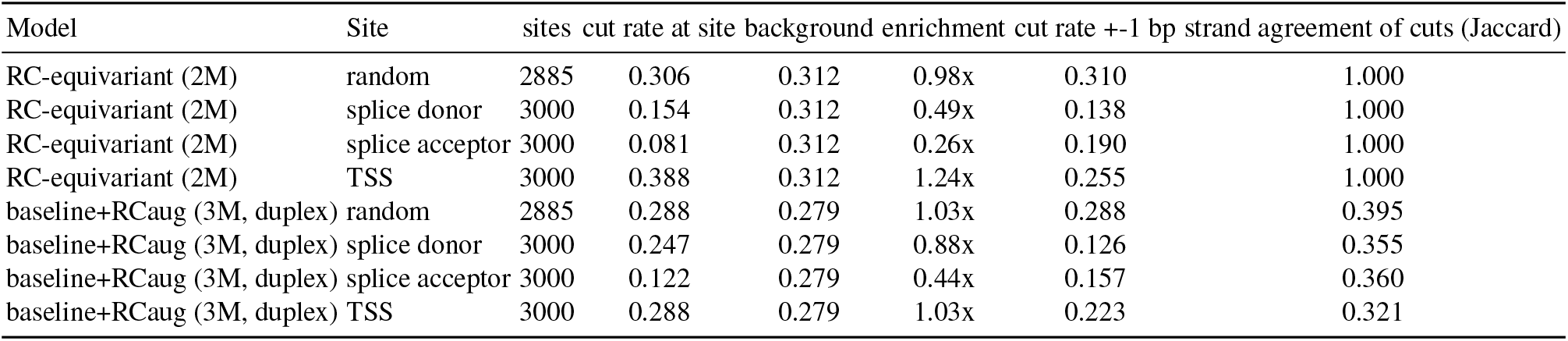
Router cut rate at gene-structure sites (RefSeq, 1-kb windows centred on the site) relative to the background rate, and the strand agreement of the cuts.

**Table S10.** Tokens formed at splice donor (3000 sites): the chunk containing the first base after the site, read on the gene’s strand, with the model’s cuts and with the same cuts circularly shifted by 100–900 bp; offset = chunk start relative to the site.

| chunk (gene strand) | share, model’s cuts | share, shifted cuts | enrichment | typical offset |
|---|---|---|---|---|
| ‘AGGT‘ | 14.8% | 1.6% | 9x | -2 |
| ‘GT‘ | 8.8% | 4.3% | 2x | 0 |
| ‘GGT‘ | 5.9% | 2.8% | 2x | -1 |
| ‘AGT‘ | 3.6% | 0.2% | 15x | -1 |
| ‘G‘ | 1.6% | 11.0% | 0x | 0 |
| ‘AGGTA‘ | 1.3% | 0.7% | 2x | -2 |
| ‘GGGT‘ | 1.1% | 0.1% | 8x | -2 |
| ‘AAGGT‘ | 1.0% | 0.5% | 2x | -3 |
| ‘TGT‘ | 1.0% | 0.2% | 4x | -1 |
| ‘AAGT‘ | 1.0% | 0.1% | 7x | -2 |
| ‘TGGT‘ | 1.0% | 0.4% | 2x | -2 |
| ‘ACAGGT‘ | 0.9% | 0.1% | 9x | -4 |

**Table S11.**
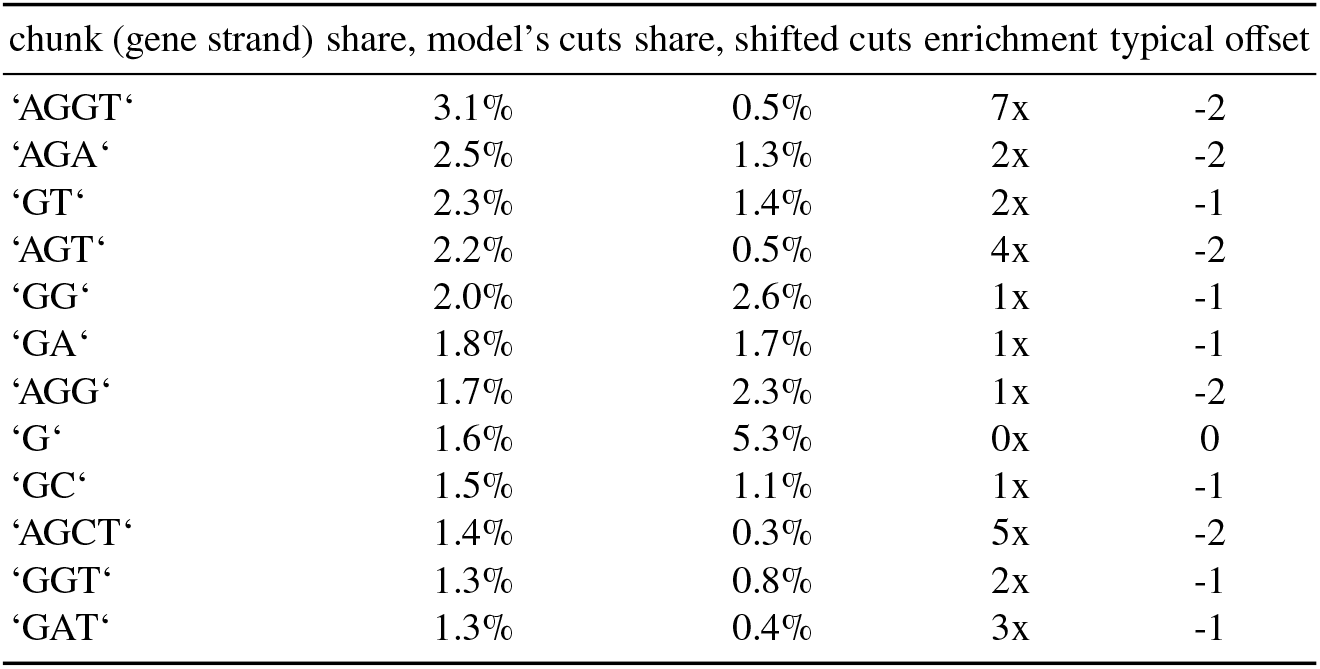
Tokens formed at splice acceptor (3000 sites): the chunk containing the first base after the site, read on the gene’s strand, with the model’s cuts and with the same cuts circularly shifted by 100–900 bp; offset = chunk start relative to the site. chunk (gene strand) share, model’s cuts share, shifted cuts enrichment typical offset

**Table S12.** Tokens formed at TSS (2127 sites): the chunk containing the first base after the site, read on the gene’s strand, with the model’s cuts and with the same cuts circularly shifted by 100–900 bp; offset = chunk start relative to the site.

| chunk (gene strand) | share, model’s cuts | share, shifted cuts | enrichment | typical offset |
|---|---|---|---|---|
| ‘A‘ | 6.4% | 5.1% | 1x | 0 |
| ‘G‘ | 1.7% | 3.3% | 1x | 0 |
| ‘GC‘ | 1.3% | 0.4% | 3x | 0 |
| ‘AGT‘ | 1.1% | 0.6% | 2x | 0 |
| ‘AGATGATG‘ | 0.9% | 0.0% | n/a | -2 |
| ‘AG‘ | 0.8% | 1.5% | 1x | 0 |
| ‘GA‘ | 0.8% | 0.6% | 1x | 0 |
| ‘C‘ | 0.8% | 0.9% | 1x | 0 |
| ‘ACT‘ | 0.8% | 0.1% | 5x | 0 |
| ‘AGC‘ | 0.8% | 0.1% | 8x | 0 |
| ‘AC‘ | 0.8% | 0.4% | 2x | 0 |
| ‘ACA‘ | 0.6% | 0.2% | 2x | -2 |

### S3.6 Accuracy with and without the outlier tasks

**Table S13.** Forward MCC averaged over the full suites and without the splice-site tasks (NT-15) and Ensembl regulatory elements (GB-8), mean _±SD_ over 3 seeds of the per-seed suite average. MirrorDNA: head chosen per task on validation. The reduced sets were chosen after inspecting the per-task results and are reported only to show where the differences are concentrated.

| Model | NT-18 | NT-15 | GB-9 | GB-8 | All 27 | All 29 |
|---|---|---|---|---|---|---|
| NT-v2 | .652<sub>±.002</sub> | .590<sub>±.002</sub> | .741<sub>±.004</sub> | .721<sub>±.005</sub> | .682<sub>±.002</sub> | .700<sub>±.002</sub> |
| DNABERT-2 | .631<sub>±.018</sub> | .587<sub>±.023</sub> | .699<sub>±.038</sub> | .684<sub>±.043</sub> | .654<sub>±.023</sub> | .674<sub>±.022</sub> |
| HyenaDNA-small | .615<sub>±.007</sub> | .564<sub>±.001</sub> | .661<sub>±.021</sub> | .667<sub>±.003</sub> | .630<sub>±.006</sub> | .645<sub>±.006</sub> |
| Caduceus-PS-1k | .595<sub>±.002</sub> | .542<sub>±.004</sub> | .725<sub>±.003</sub> | .714<sub>±.008</sub> | .638<sub>±.002</sub> | .595<sub>±.004</sub> |
| Caduceus-PH-1k | .579<sub>±.003</sub> | .532<sub>±.003</sub> | .713<sub>±.005</sub> | .698<sub>±.007</sub> | .624<sub>±.001</sub> | .640<sub>±.002</sub> |
| Chunker + RC aug. | .587<sub>±.005</sub> | .553<sub>±.005</sub> | .706<sub>±.005</sub> | .723<sub>±.003</sub> | .627<sub>±.005</sub> | .645<sub>±.005</sub> |
| MirrorDNA-duplex | .583<sub>±.002</sub> | .541<sub>±.002</sub> | .738<sub>±.003</sub> | .746<sub>±.001</sub> | .635<sub>±.002</sub> | .655<sub>±.002</sub> |
| MirrorDNA (ours) | .618<sub>±.003</sub> | .549<sub>±.004</sub> | .751<sub>±.002</sub> | .750<sub>±.004</sub> | .662<sub>±.003</sub> | .682<sub>±.003</sub> |

**Table S14.** MirrorDNA against each model: tasks on which MirrorDNA has the higher mean MCC over 3 seeds / tasks (first row) and two-sided Wilcoxon signed-rank *p* over tasks (second row). Dir.: the 6 direction-dependent NT tasks; Gene: the 2 gene-strand tasks (too few for a test).

| MirrorDNA vs. | NT-18 | NT-15 | Dir. | GB-9 | GB-8 | Gene | All 27 | All 29 |
|---|---|---|---|---|---|---|---|---|
| NT-v2 | 3/18<br>(.001) | 1/15<br>(<.001) | 3/6<br>(.84) | 6/9<br>(.36) | 6/8<br>(.078) | 1/2<br>– | 9/27<br>(.055) | 10/29<br>(.076) |
| DNABERT-2 | 4/18<br>(.42) | 1/15<br>(.008) | 3/6<br>(.44) | 5/9<br>(.82) | 5/8<br>(.55) | 2/2<br>– | 9/27<br>(.53) | 11/29<br>(.56) |
| HyenaDNA-small | 8/18<br>(.64) | 5/15<br>(.048) | 6/6<br>(.031) | 9/9<br>(.004) | 8/8<br>(.008) | 2/2<br>– | 17/27<br>(.052) | 19/29<br>(.029) |
| Caduceus-PS-1k | 14/18<br>(.005) | 11/15<br>(.035) | 6/6<br>(.031) | 7/9<br>(.20) | 7/8<br>(.078) | 2/2<br>– | 21/27<br>(.004) | 23/29<br>(<.001) |
| Caduceus-PH-1k | 14/18<br>(.008) | 11/15<br>(.055) | 6/6<br>(.031) | 7/9<br>(.16) | 7/8<br>(.039) | 2/2<br>– | 21/27<br>(.003) | 23/29<br>(<.001) |
| Chunker + RC aug. | 8/18<br>(.93) | 5/15<br>(.21) | 5/6<br>(.062) | 6/9<br>(.30) | 5/8<br>(.55) | 2/2<br>– | 14/27<br>(.47) | 16/29<br>(.21) |
| MirrorDNA-duplex | 12/18<br>(.018) | 9/15<br>(.12) | 6/6<br>(.031) | 6/9<br>(.30) | 5/8<br>(.55) | 2/2<br>– | 18/27<br>(.008) | 20/29<br>(.002) |

### S3.7 Frozen representations on the two strands

**Table S15.** Frozen pretrained models on the 12 strand-symmetric NT tasks (mean over tasks; no fine-tuning). cos: cosine similarity between the standardized pooled embeddings of a test sequence and of its reverse complement; nn: share of reverse-complemented test sequences whose nearest neighbour among the forward test embeddings is the same DNA (values slightly below 1 for the equivariant models come from near-duplicate test sequences); a logistic-regression probe is trained on forward embeddings (up to 20,000 sequences per task) and evaluated on the test set as given (fwd) and reverse-complemented (RC), with the both-strands MCC, the flip rate, and the MCC of the probe averaged over both strands (avg.). Cosine values are computed in each model’s own embedding space and are best compared within a model.

| Model | cos | nn | fwd | RC | both | flips | avg. |
|---|---|---|---|---|---|---|---|
| MirrorDNA (ours) | 1.000 | 0.992 | 0.449 | 0.449 | **0.449** | 0.0% | 0.449 |
| Caduceus-PS-1k | 1.000 | 0.992 | 0.441 | 0.441 | 0.441 | 0.0% | 0.441 |
| Caduceus-PH-1k | 0.582 | 0.041 | 0.436 | 0.398 | 0.273 | 14.4% | 0.430 |
| DNABERT-2 | 0.215 | 0.053 | 0.447 | 0.412 | 0.207 | 22.5% | 0.464 |
| NT-v2 | 0.223 | 0.023 | 0.443 | 0.380 | 0.193 | 22.0% | 0.446 |
| Chunker + RC aug. | 0.588 | 0.098 | 0.476 | 0.336 | 0.148 | 25.8% | 0.438 |
| HyenaDNA-small | 0.587 | 0.028 | 0.464 | 0.351 | 0.174 | 23.4% | 0.445 |

### S3.8 Both strands, strand-symmetric tasks

**Table S16.** Fine-tuned models on the 12 strand-symmetric NT tasks and the 7 strand-symmetric Genomic Benchmarks tasks, evaluated on the test sets as given (fwd) and reverse-complemented (RC): MCC, both-strands MCC and flip rate (%), mean _±SD_ over 3 seeds of the mean over tasks. The two direction-dependent GB tasks (coding vs. intergenomic, non-TATA promoters) are excluded. MirrorDNA: head chosen per task on validation.

| Model | symmetric NT (12) |  |  |  | symmetric GB (7) |  |  |  |
|---|---|---|---|---|---|---|---|---|
|  | fwd | RC | both | flips | fwd | RC | both | flips |
| NT-v2 | .539<sub>±.004</sub> | .334<sub>±.006</sub> | .152<sub>±.009</sub> | 28.2<sub>±0.5</sub> | .710<sub>±.006</sub> | .685<sub>±.003</sub> | .588<sub>±.007</sub> | 11.1<sub>±0.4</sub> |
| DNABERT-2 | .537<sub>±.030</sub> | .350<sub>±.012</sub> | .187<sub>±.006</sub> | 25.4<sub>±2.1</sub> | .643<sub>±.048</sub> | .620<sub>±.051</sub> | .549<sub>±.056</sub> | 8.4<sub>±0.7</sub> |
| HyenaDNA-small | .515<sub>±.001</sub> | .272<sub>±.004</sub> | .080<sub>±.004</sub> | 31.1<sub>±0.3</sub> | .616<sub>±.026</sub> | .595<sub>±.027</sub> | .466<sub>±.029</sub> | 14.2<sub>±0.6</sub> |
| Caduceus-PS-1k | .487<sub>±.002</sub> | .487<sub>±.002</sub> | .487<sub>±.002</sub> | 0.0<sub>±0.0</sub> | .686<sub>±.005</sub> | .686<sub>±.005</sub> | .686<sub>±.005</sub> | 0.0<sub>±0.0</sub> |
| Caduceus-PH-1k | .489<sub>±.004</sub> | .486<sub>±.003</sub> | .389<sub>±.006</sub> | 9.8<sub>±0.7</sub> | .679<sub>±.007</sub> | .677<sub>±.006</sub> | .595<sub>±.009</sub> | 8.3<sub>±1.5</sub> |
| Chunker + RC aug. | .498<sub>±.006</sub> | .484<sub>±.006</sub> | .381<sub>±.007</sub> | 11.0<sub>±0.2</sub> | .652<sub>±.006</sub> | .655<sub>±.014</sub> | .561<sub>±.015</sub> | 9.8<sub>±1.4</sub> |
| MirrorDNA-duplex | .487<sub>±.002</sub> | .487<sub>±.002</sub> | .487<sub>±.002</sub> | 0.0<sub>±0.0</sub> | .693<sub>±.004</sub> | .693<sub>±.004</sub> | .693<sub>±.004</sub> | 0.0<sub>±0.0</sub> |
| MirrorDNA (ours) | .491<sub>±.004</sub> | .491<sub>±.004</sub> | .491<sub>±.004</sub> | 0.0<sub>±0.0</sub> | .711<sub>±.002</sub> | .711<sub>±.002</sub> | .711<sub>±.002</sub> | 0.0<sub>±0.0</sub> |

## S4 Additional figures

**Fig. S1.**
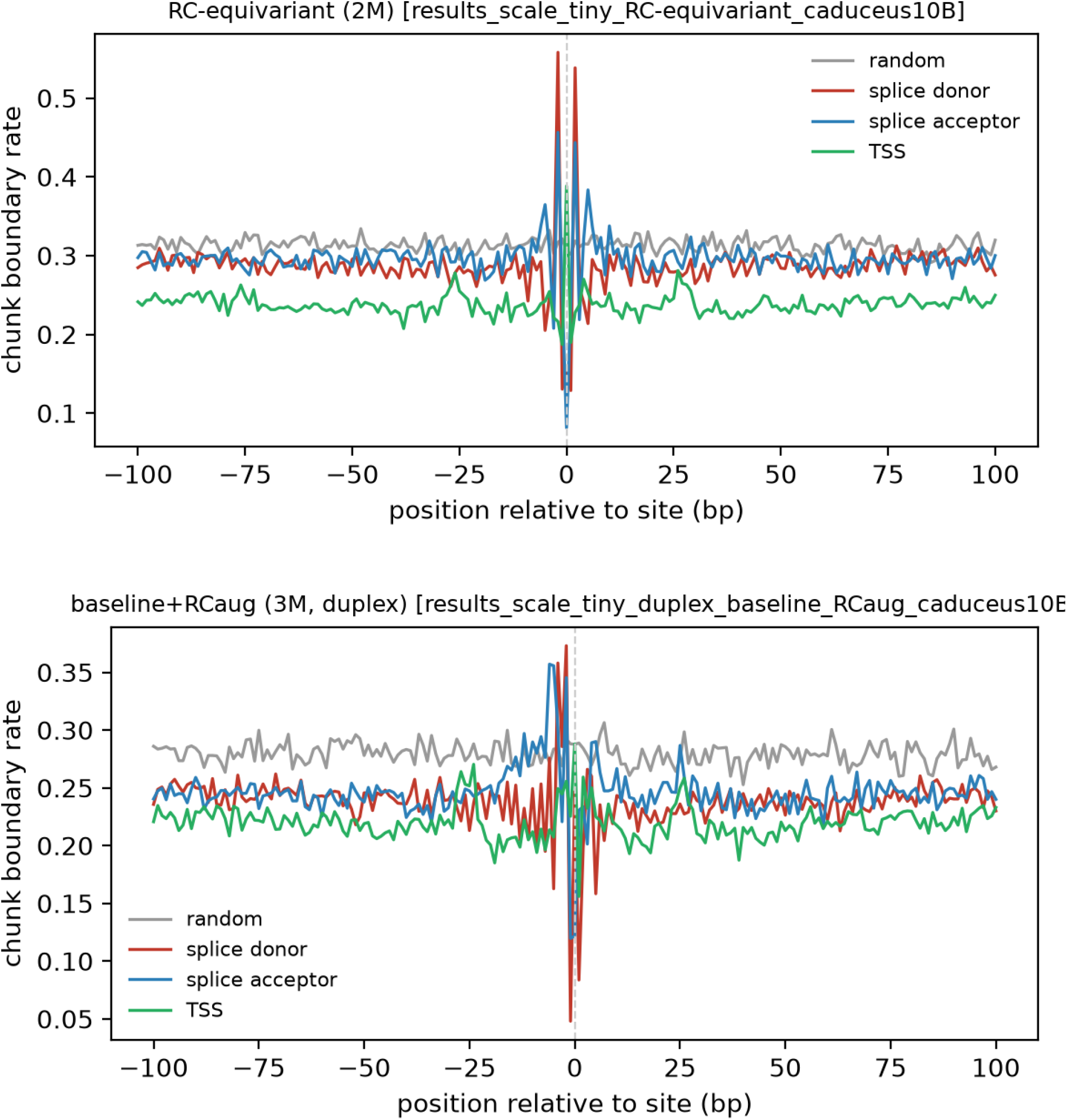
Router cut rate around gene-structure sites relative to the background rate: MirrorDNA (top) and the augmented chunker (bottom).

**Fig. S2.**
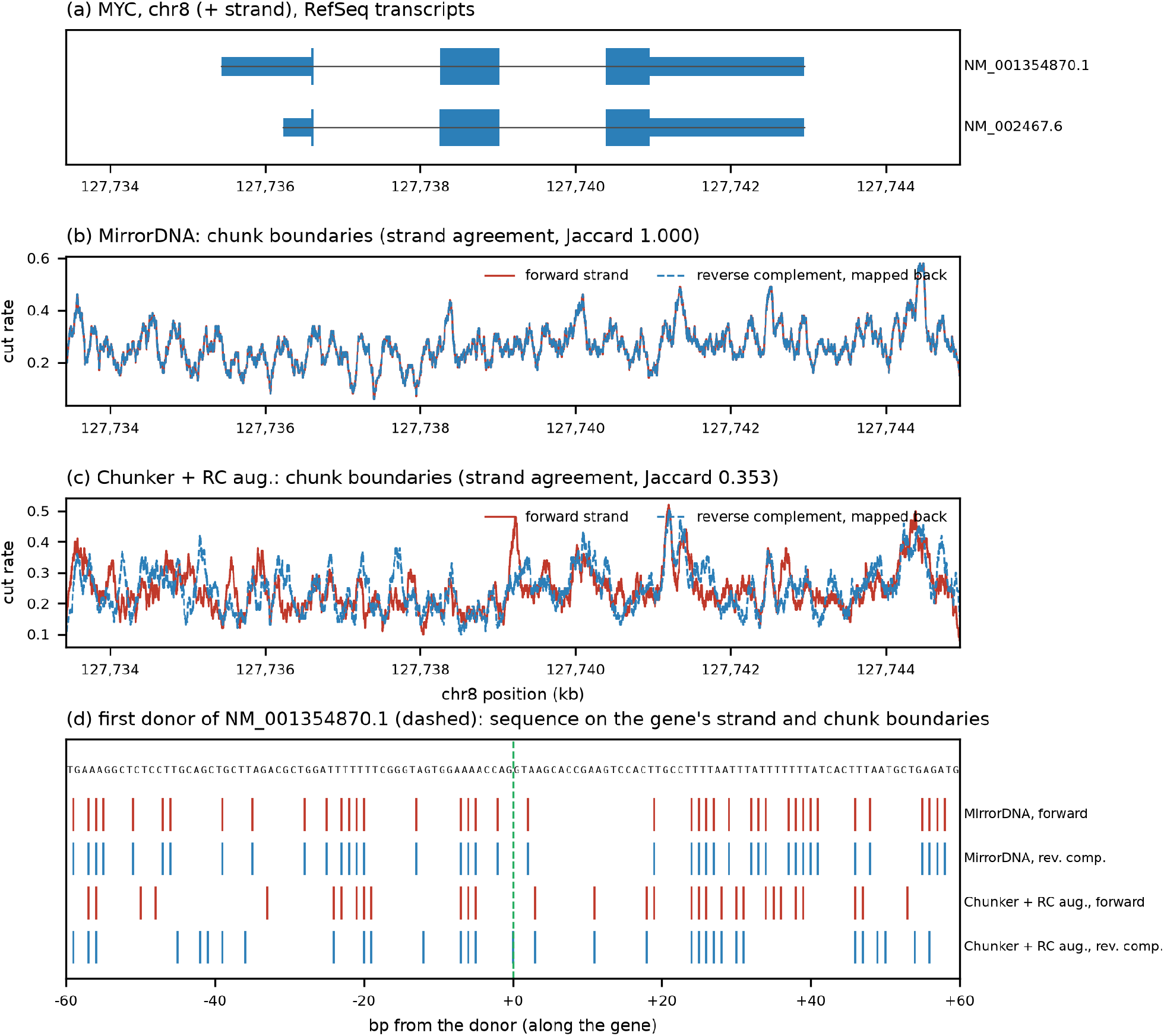
**MYC** (chr8, plus strand). Panels as in Fig. 6.

**Table S17.**
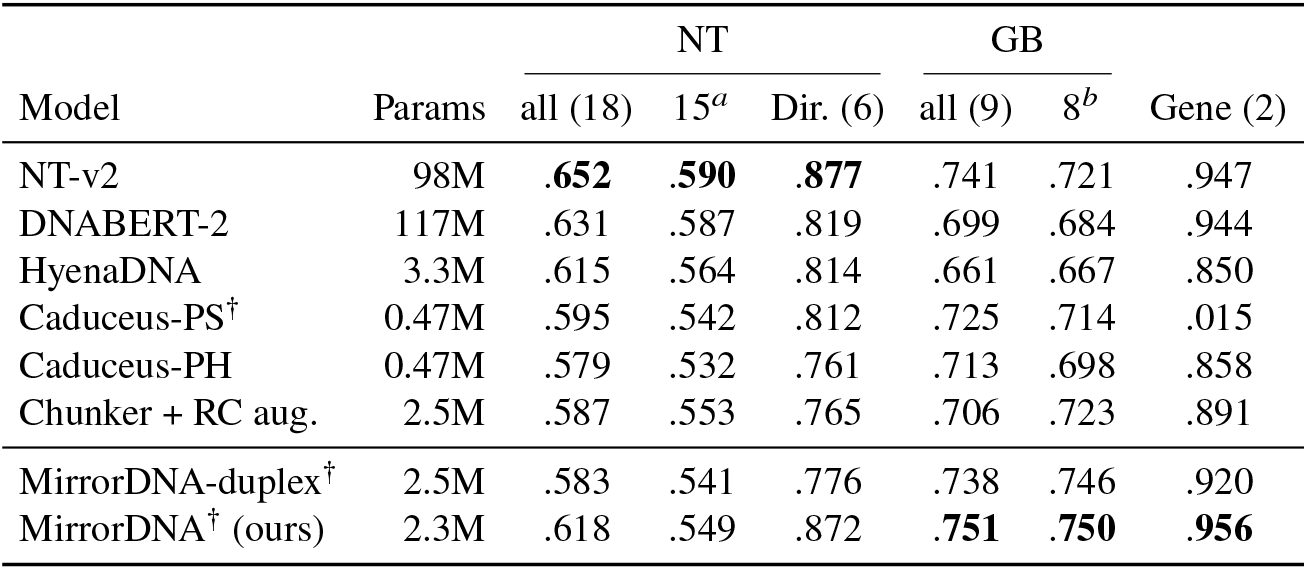
Accuracy. (forward MCC, mean over 3 seeds) on the Nucleotide Transformer (NT) tasks, the Genomic Benchmarks (GB) and the 2 gene-strand tasks (Gene). Dir. = the 6 direction-dependent NT tasks. ^*a*^ NT without the 3 splice-site tasks, which depend on single-base precision at the junction. ^*b*^ GB without Ensembl regulatory elements, on which all three chunk-based models trail the single-nucleotide models. MirrorDNA: head chosen per task on validation (Section 2.4). Bold: best per column. Caduceus: the 1-kb checkpoints. SDs: Supplementary Table S13; per-task results: Supplementary Section S3. ^†^Exactly equivariant.

|  |  | NT |  |  | GB |  |  |
|---|---|---|---|---|---|---|---|
| Model | Params | all (18) | 15<sup>*a*</sup> | Dir. (6) | all (9) | 8<sup>*b*</sup> | Gene (2) |
| NT-v2 | 98M | .**652** | .**590** | .**877** | .741 | .721 | .947 |
| DNABERT-2 | 117M | .631 | .587 | .819 | .699 | .684 | .944 |
| HyenaDNA | 3.3M | .615 | .564 | .814 | .661 | .667 | .850 |
| Caduceus-PS<sup>†</sup> | 0.47M | .595 | .542 | .812 | .725 | .714 | .015 |
| Caduceus-PH | 0.47M | .579 | .532 | .761 | .713 | .698 | .858 |
| Chunker + RC aug. | 2.5M | .587 | .553 | .765 | .706 | .723 | .891 |
| MirrorDNA-duplex<sup>†</sup> | 2.5M | .583 | .541 | .776 | .738 | .746 | .920 |
| MirrorDNA<sup>†</sup> (ours) | 2.3M | .618 | .549 | .872 | .**751** | .**750** | .**956** |

## Notes

### Competing Interest Statement

The authors have declared no competing interest.

